# T-type Ca^2+^ and persistent Na^+^ currents synergistically elevate ventral, not dorsal, entorhinal cortical stellate cell excitability

**DOI:** 10.1101/2021.05.07.443063

**Authors:** Aleksandra Topczewska, Elisabetta Giacalone, Wendy S. Pratt, Michele Migliore, Annette C. Dolphin, Mala M. Shah

**Author notes:** Corresponding author: Mala M. Shah, Department of Pharmacology, UCL School of Pharmacy, 29-39 Brunswick Square, London, WC1N 1AX, UK.

## Abstract

The medial entorhinal cortex (mEC) plays a salient role in physiological processes such as spatial cognition and spatial coding. mEC layer II stellate neurons, in particular, influence these processes. Interestingly, ventral and dorsal stellate neurons diversely affect these processes and have distinct intrinsic membrane properties and action potential firing patterns. Little, though, is known about how ventral stellate neuron intrinsic excitability is regulated. We show that ventral stellate neurons predominantly possess T-type Ca^2+^ currents encoded by Ca_V_3.2 subunits, with dorsal stellate neurons having small or no currents. Further, twice as much Ca_V_3.2 mRNA was present in ventral than dorsal mEC. In line with T-type, Ca_V_3.2 Ca^2+^ current biophysical properties, depolarising stimuli activated these currents in ventral, but not dorsal, neurons. Here, these currents acted in concert with persistent Na^+^ currents to elevate input resistance and tonic action potential firing. Ca_V_3.2 currents also enhanced excitatory post-synaptic potential decay and integration solely in ventral neurons. These results reveal that Ca_V_3.2 currents, together with persistent Na^+^ currents, impart the characteristic intrinsic membrane and firing properties of ventral stellate neurons. This signifies that specific voltage-gated conductances distinctly affect ventral and dorsal mEC stellate neuron activity and functions such as spatial memory and spatial navigation.

---

The medial entorhinal cortex (mEC) forms a significant component of the hippocampal formation. It is established that the mEC critically influences episodic memory formation, spatial memory and spatial coding (Hardcastle et al., 2017; Hasselmo et al., 2009; Rowland et al., 2016; Sasaki et al., 2015). Interestingly, dorsal mEC neurons influence spatial memory and spatial coding preferentially (Steffenach et al., 2005; Strange et al., 2014). In contrast, ventral mEC lesions reduced defensive behaviour on an elevated plus maze, indicating that these neurons predominantly affect emotional behaviour (Steffenach et al., 2005). This accords with the prevailing view that dorsal and ventral mEC neurons have distinctive functions in influencing physiological processes such as cognition (Cembrowski and Spruston, 2019; Fanselow and Dong, 2010; Strange et al., 2014).

Consistent with this, emerging evidence indicates that dorsal and ventral mEC neurons form unique circuit connections with other cortical and subcortical regions (Strange et al., 2014). In keeping with this, dorsal and ventral mEC layer II and III neurons predominantly connect to dorsal and ventral hippocampal dentate gyrus granule and pyramidal neurons respectively (Strange et al., 2014). Importantly, recent evidence indicates that gene expression is distinct in ventral and dorsal mEC and hippocampal neurons and their downstream targets (Cembrowski and Spruston, 2019; Gergues et al., 2020; Papatheodoropoulos, 2018; Thompson et al., 2008). Accordingly, ion channel densities and biophysical properties, intrinsic membrane properties as well as intrinsic and synaptic excitability differ along the dorsal-ventral axis within the mEC (Bant et al., 2020; Beed et al., 2013; Garden et al., 2008; Giocomo and Hasselmo, 2008, 2009; Pastoll et al., 2012a). This has been predominantly characterised in mEC layer II stellate neurons. Indeed, dorsal stellate neurons have higher densities and altered biophysical properties of K^+^ channels and the hyperpolarization-activated cyclic nucleotide-gated (HCN) channels compared with their ventral neuron counterparts (Garden et al., 2008; Giocomo and Hasselmo, 2008, 2009; Pastoll et al., 2012a).

This has been suggested to render dorsal stellate neurons less excitable than their ventral equivalents. It is noteworthy, though, that dorsal stellate neurons have a significantly larger density of subthreshold Na^+^ currents than ventral stellate neurons (Bant et al., 2020). This, therefore, raises the question of how ventral stellate neuron excitability is maintained at higher levels than dorsal neurons. In particular, it remains to be determined whether there are specific voltage-gated conductances that significantly influence ventral stellate neuron intrinsic and synaptic excitability.

We explored whether T-type Ca^2+^ channels might regulate ventral and/or dorsal stellate neuron activity distinctly as these channels are subthreshold-active voltage-gated ion channels that significantly contribute to shaping neuronal firing patterns. These channels have been shown to promote low threshold Ca^2+^ spikes and so-called ‘burst firing’ (i.e. a brief clustering of action potentials) (Cain and Snutch, 2010; Crunelli et al., 2018; Leresche and Lambert, 2017; Weiss and Zamponi, 2019; Zamponi et al., 2015). Since hyperpolarizing potentials such as inhibitory postsynaptic potentials enable these channels to recover from inactivation, they also facilitate rebound action potential firing following termination of brief hyperpolarizing potentials in some neurons (Cain and Snutch, 2010; Weiss and Zamponi, 2019; Zamponi et al., 2015). Further, recent evidence suggests that these channels modify synaptic release and boost excitatory post-synaptic potential (EPSP) integration in some neurons, thereby contributing to synaptic plasticity (Leresche and Lambert, 2017).

Interestingly, low voltage-activated Ca^2+^ currents with properties similar to T-type Ca^2+^ currents have been identified in isolated adult mEC layer II stellate neurons (Bruehl and Wadman, 1999). It is, however, unknown if there is a gradient in these currents along the dorsal-ventral mEC axis and how these currents affect mEC dorsal/ventral neuron excitability.

Here, we report that T-type Ca^2+^ currents were predominantly present in ventral stellate neurons only. Moreover, expression of the T-type Ca^2+^ channel α1 subunit, Ca_V_3.2, in ventral mEC was twice that in dorsal mEC. The T-type, Ca_V_3.2 Ca^2+^ currents were activated by depolarizing potentials in mEC ventral stellate neurons only. Accordingly, these currents prolonged EPSPs and enhanced EPSP summation in these neurons, suggesting that they selectively impact synaptic potential information processing here. Additionally, and unusually, these currents acted in concert with subthreshold Na^+^ currents to specifically augment ventral stellate neuron input resistance and, consequently, tonic action potential firing. This is a previously undescribed cellular mechanism by which T-type Ca^2+^ channels raise excitability in central neurons. These findings also strongly support the notion that ventral and dorsal mEC stellate neurons express distinct voltage-dependent conductances that regulate their activity and which might influence their discrete roles in processes such as cognition.

## Results

### T-type, Ca_V_3.2 Ca^2+^ currents enhance wildtype ventral, not dorsal, stellate neuron action potential firing propensity

T-type Ca^2+^ channels are encoded by 3 α1 subunits: Ca_V_3.1, Ca_V_3.2 and Ca_V_3.3. We have previously shown that Ca_V_3.2 is predominantly expressed in the mEC(Huang et al., 2011). Thus, to determine how T-type Ca^2+^ currents influence mEC layer II stellate neuron activity, we made electrophysiological recordings from these neurons present in parasagittal brain slices obtained from adult wildtype and Ca_V_3.2 null (Ca_V_3.2^-/-^) littermate mice in the presence of glutamate and GABA receptor inhibitors (see Materials and Methods). Neurons were identified as being dorsal or ventral if they were located within 0 % - 30 % or 70 % - 100 % of the mEC - parasubiculum border respectively (Pastoll et al., 2012b) (Fig 1a, 1b; see Materials and Methods). Stellate neurons were visually identified as having a polygonal or ovoid soma with multiple dendrites radiating from this (Fig 1a, 1b; Supp Fig 1a). All patched neurons were filled with neurobiotin and subsequently stained with Alexa Fluor 488- streptavidin coupled antibodies (see Materials and Methods). Full morphological analysis was carried out from a subset of stained neurons (Supp Fig 1). Sholl analysis revealed no differences in dendrite numbers between dorsal and ventral wildtype and Ca_V_3.2^-/-^ stellate neurons (Supp Fig 1b). Consistent with previous reports (Garden et al., 2008), stellate neuron cell body area, though, was greater in dorsal than ventral locations in wildtype and Ca_V_3.2^-/-^ slices (Supp Fig 1c).

**Fig 1:**
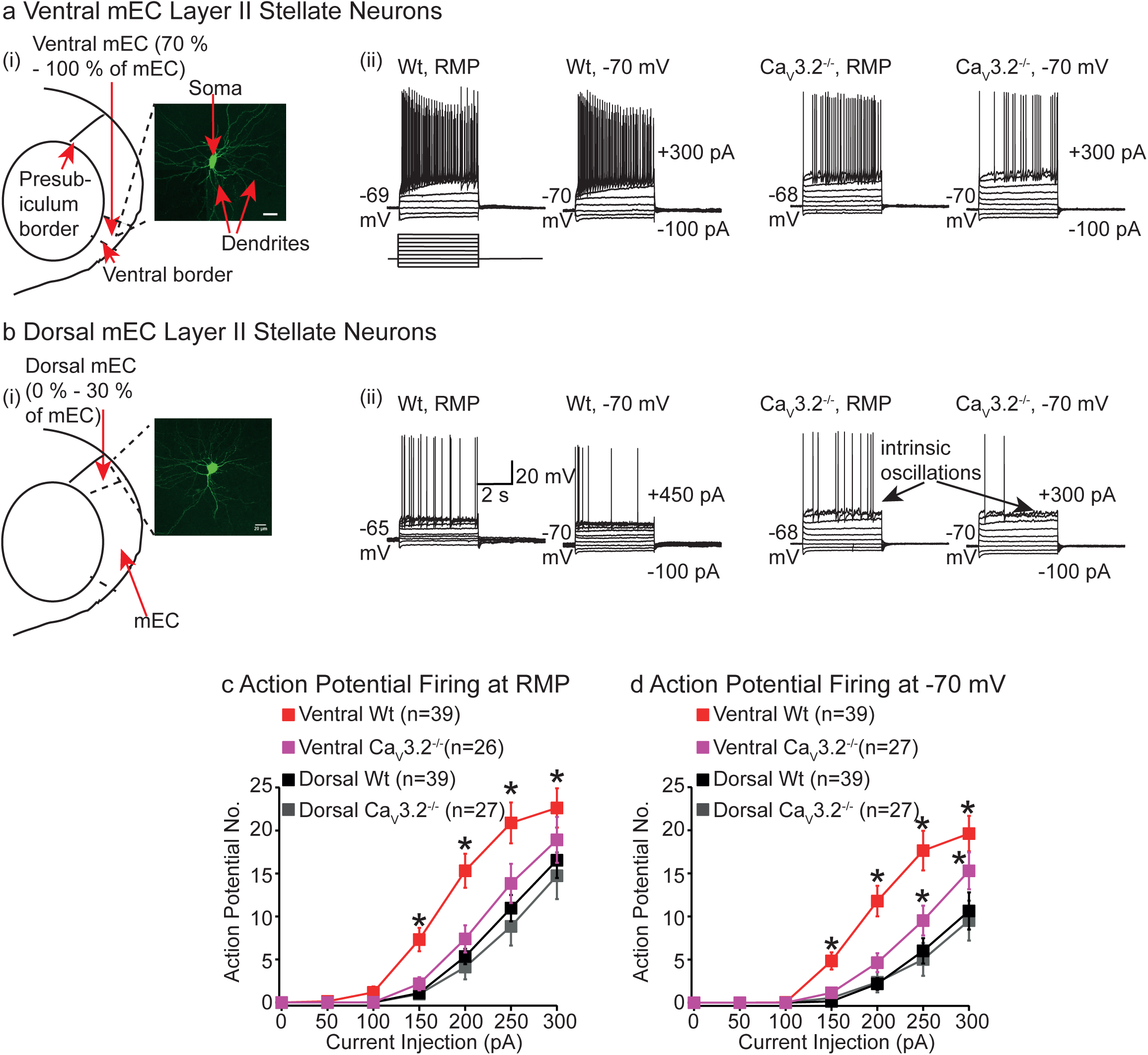
Ca_V_3.2 Ca^2+^ currents raise ventral, not dorsal, layer II stellate neuron excitability. **a(i), b(i)** Schematics illustrating ventral and dorsal mEC regions respectively. Also shown are representative confocal images of a dorsal and ventral mEC layer II stellate neurons. The scale on both images depicts 20 μm. **a(ii), b(ii)** Typical whole-cell current- clamp recordings from ventral and dorsal mEC wildtype (Wt) and Ca_V_3.2-^/-^ layer II stellate neurons respectively either at the resting membrane potential (RMP) or at a fixed potential of -70 mV when 5 s long hyperpolarizing and depolarizing steps from -100 pA in increments of 50 pA were applied. The membrane potential is indicated adjacent to each trace. In addition for each set of traces, the current injections applied to elicit the recordings are shown next to the -70 mV traces. The scale associated with the first panel applies to all panels. **c, d** Plots showing the numbers of action potentials elicited by 5 s long, depolarising steps of varying current amplitude at RMP and -70 mV respectively. **c,** * signifies p < 0.05 when compared between all other groups using a two-way ANOVA with Fisher’s Least Significance Difference (LSD) post hoc test (exact p values are stated in Supp Table 7). **d,** Asterisk for ventral wildtype group indicates significance at p < 0.05 when compared with all other groups whereas the asterisk associated with ventral Ca_V_3.2-/- group depicts significance when compared with dorsal wildtype and Ca_V_3.2^-/-^ groups. All comparisons were made using a two-way ANOVA with Fisher’s LSD post hoc test. Exact p values are in Supp Table 7. The numbers of observations for each group are shown in parenthesis.

Interestingly, whole-cell current clamp recordings revealed that whilst abolition of Ca_V_3.2 subunit expression had little effect on ventral mEC layer II stellate neuron resting membrane potential (RMP; Supp Table 1), the numbers of action potentials generated with 5 s long, depolarizing steps either at RMP or a fixed potential of -70 mV were significantly fewer in ventral Ca_V_3.2^-/-^ neurons than ventral wildtypes (Fig 1a, 1c, 1d). The time taken for the first action potential to be initiated, though, with the smallest depolarizing step was similar between the two groups (Supp Table 1). There were also no differences in action potential firing patterns between ventral wildtype and Ca_V_3.2^-/-^ neurons (Fig 1a, data not shown).

Moreover, the action potential threshold and shape measured by eliciting single action potentials with short, 10 ms pulses at a fixed potential of – 70 mV were unaffected by the absence of Ca_V_3.2 Ca^2+^ channels (Supp Fig 2). Further, consistent with previous studies indicating that voltage-gated Ca^2+^ channels have little effect on the generation of subthreshold oscillations in mEC stellate neurons (Klink and Alonso, 1993), similar numbers of ventral wildtype (33 % (13/39)) and Ca_V_3.2^-/-^ neurons (44 % (13/27)) displayed subthreshold oscillations elicited with depolarizing stimuli (data not shown). Hence, Ca_V_3.2 Ca^2+^ channels have a profound effect on the propensity for action potentials to be elicited with depolarizing stimuli in ventral wildtype stellate neurons without affecting the RMP or action potential threshold.

The lack of Ca_V_3.2 subunit expression, though, had no effect on dorsal mEC layer II stellate RMP (Supp Table 1), intrinsic oscillations (data not shown), action potential numbers elicited with depolarizing pulses either at RMP or a fixed potential of -70 mV (Fig 1b, 1c, 1d) and action potential threshold (Supp Fig 2). This indicates that Ca_V_3.2 subunit expression varies along the dorsal ventral mEC axis and influences ventral neuron excitability predominantly. Indeed, whilst ventral wildtype neurons were significantly more excitable than dorsal wildtype neurons (Fig 1), similar numbers of action potentials were elicited in ventral and dorsal Ca_V_3.2^-/-^ stellate neurons at RMP (Fig 1a, 1c). Further, at a fixed potential of -70 mV, comparable numbers of spikes were produced with small amplitude (< 200 pA) - depolarizing pulses in ventral and dorsal Ca_V_3.2^-/-^ neurons (Fig 1d). These findings, thus, strongly suggest that T-type, Ca_V_3.2 Ca^2+^ channels play a significant role in influencing mEC layer II stellate neuron spike firing across the dorsal-ventral axis by preferentially enhancing action potential rates in ventral wildtype neurons alone.

### T-type Ca^2+^ channel inhibitors selectively reduce ventral wildtype stellate neuron action potential firing rates

To further investigate the notion that T-type Ca^2+^ channels solely enhance ventral wildtype stellate neuron activity, we tested the effects of the specific T-type Ca^2+^ channel inhibitors, TTA-P2 (100 nM(Shipe et al., 2008)) and NiCl_2_ (50 μM, a concentration that preferentially inhibits Ca_V_3.2 Ca^2+^ channels (Lee et al., 1999)) on ventral and dorsal wildtype and Ca_V_3.2^-/-^ stellate neurons. 20 min external application of TTA-P2 or NiCl_2_ had little effect on dorsal and ventral wildtype and Ca_V_3.2^-/-^ stellate neuron RMP (Supp Table 2), confirming that T-type Ca^2+^ channels are not active at RMP.

Bath application of TTA-P2 (Fig 2) or NiCl_2_ (Supp Fig 3) onto ventral wildtype stellate neurons, though, significantly inhibited the number of action potentials induced by depolarising pulses either at a fixed potential of -70 mV (Fig 2a(i), 2a(ii), Supp Fig 3a) or at the normal RMP (data not shown). This effect was partially reversible upon 20 min washout of NiCl_2_ (Supp Fig 3), but not TTA-P2 (Fig 2a(i), 2a(ii)). This accords with the slow reversal of the effects of TTA-P2 on T-type Ca^2+^ current in other neuron subtypes (Choe et al., 2011; Dreyfus et al., 2010). Again, T-type Ca^2+^ channel inhibitors had little effect on action potential threshold, amplitude and half-width of single action potentials at -70 mV on ventral wildtype neurons (Supp Table 3), indicating that T-type Ca^2+^ channels significantly alter ventral wildtype neuron spike firing rates without affecting action potential characteristics or the RMP.

**Fig 2:**
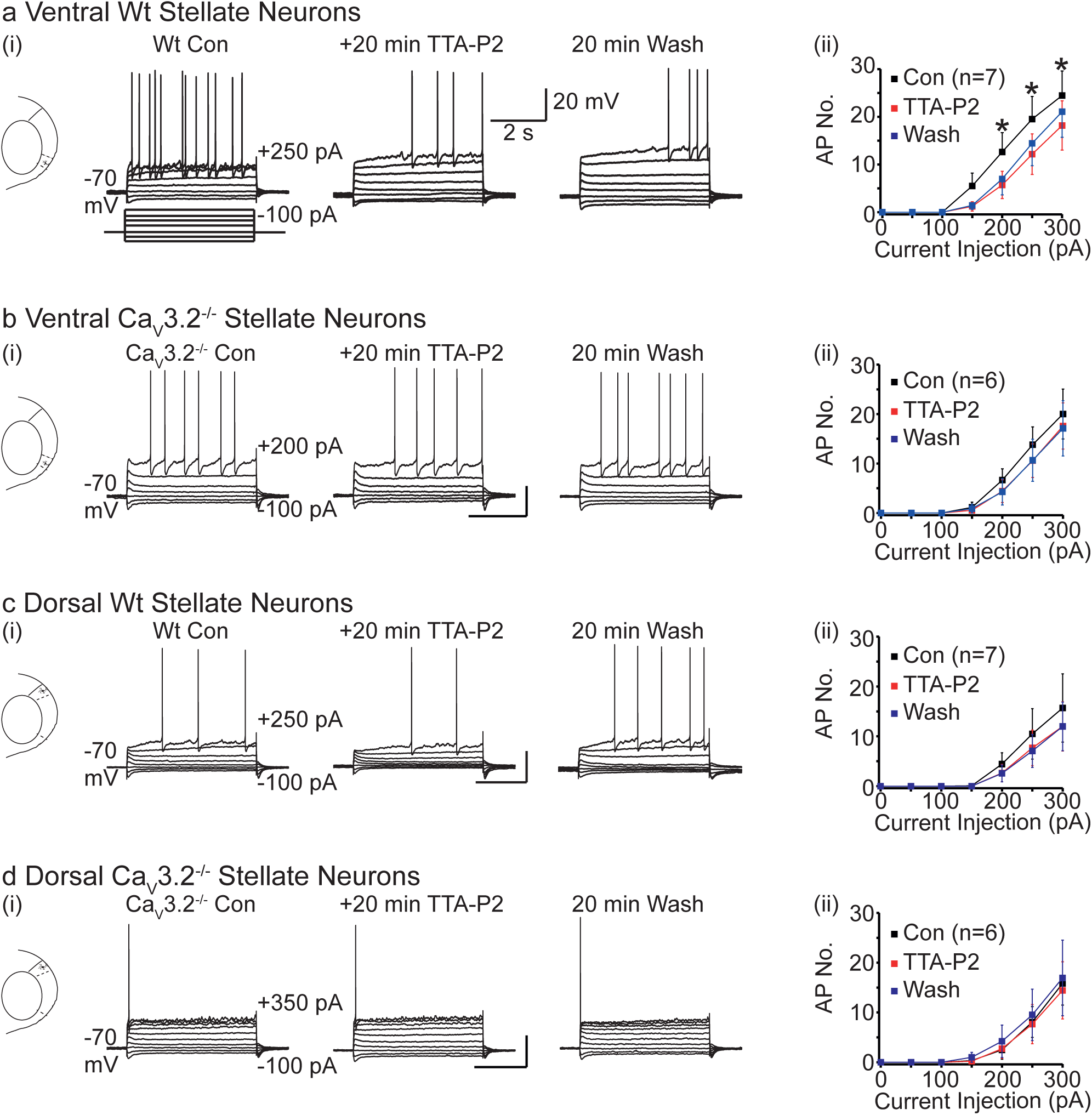
TTA-P2 inhibits action potential firing solely in ventral wildtype stellate neurons. **a(i), b(i), c(i), d(i)** Representative traces from ventral wildtype, ventral Ca_V_3.2^-/-^, dorsal wildtype and dorsal Ca_V_3.2^-/-^ stellate neurons respectively under control (Con) conditions, following 20 min application of 100 nM TTA-P2 and after 20 min washout (wash) of TTA- P2. The recordings were generated by applying 5 s long, 50 pA incremental steps at a fixed potential of -70 mV as shown in the schematic in **a(i)**. The specific current injections applied to generate the recordings are stated next to the first trace in each panel. The x and y ordinates of the scale associated with each middle trace denotes 20 mV and 2 s respectively and applies to all traces in the panel. **a(ii), b(ii), c(ii), d(ii)** Graphs show numbers of action potentials elicited by applying current injections of varying amplitude in the four groups of neurons in the absence (con), presence and following washout of TTA-P2. The numbers of observations are shown in parenthesis. Asterisks signify significance at p < 0.05 when paired t-tests were applied to compare action potentials generated under control conditions and following TTA-P2 application (exact p values are stated in Supp Table 7).

In contrast, the number of spikes generated by depolarizing pulses was unaffected by either TTA-P2 or NiCl_2_ in ventral Ca_V_3.2^-/-^, dorsal wildtype or dorsal Ca_V_3.2^-/-^ stellate neurons (Fig 2b, 2c, 2d, Supp Fig 3). These inhibitors also had little effect on spike properties in ventral Ca_V_3.2^-/-^, dorsal wildtype or dorsal Ca_V_3.2^-/-^ stellate neurons (Supp Table 3). These findings substantiate the concept that Ca_V_3.2 Ca^2+^ currents promote tonic action potential firing in ventral, and not dorsal, mEC stellate neurons.

### Ca_V_3.2 currents mask the sag and increase R_N_ measured using depolarizing potentials in wildtype ventral stellate neurons alone

Next, we asked how might T-type, Ca_V_3.2 channels enhance action potential firing in ventral layer II stellate neurons, given that there were no differences in RMP, action potential threshold or spike shape between wildtype and Ca_V_3.2^-/-^ neurons and T-type Ca^2+^ channel inhibitors had little effect on these characteristics in ventral wildtype neurons (Supp Table 1, Supp Fig 2, Supp Table 2, Supp Table 3). Increases in input resistance (R_N_) are known to enhance action potential firing in stellate neurons (Garden et al., 2008; Pastoll et al., 2012a). We, thus, explored whether T-type Ca^2+^ currents affect R_N_ in ventral and dorsal neurons.

Since Ca_V_3.2 Ca^2+^ channels are activated by depolarization, it was not surprising that there were no differences in R_N_ between ventral wildtype and Ca_V_3.2^-/-^ neurons when measured using hyperpolarizing, -100 pA, 5 s step at either a fixed potential of -70 mV or at normal RMP (Supp Table 1). In contrast, when measured using depolarizing +100 pA 5 s steps at - 70 mV or normal RMP, R_N_ was significantly greater in ventral wildtype neurons compared with ventral Ca_V_3.2^-/-^ neurons (Fig 3a(i), Fig 3a(ii), Supp Table 1). The greater R_N_ in ventral wildtype neurons was due to a sustained depolarization induced by +100 pA current pulses (Fig 3a(i)), which was absent in Ca_V_3.2^-/-^ neurons. This was unexpected given that Ca_V_3.2 subunits, when expressed in heterologous systems, result in fast inactivating currents (Chemin et al., 2002).

**Fig 3:**
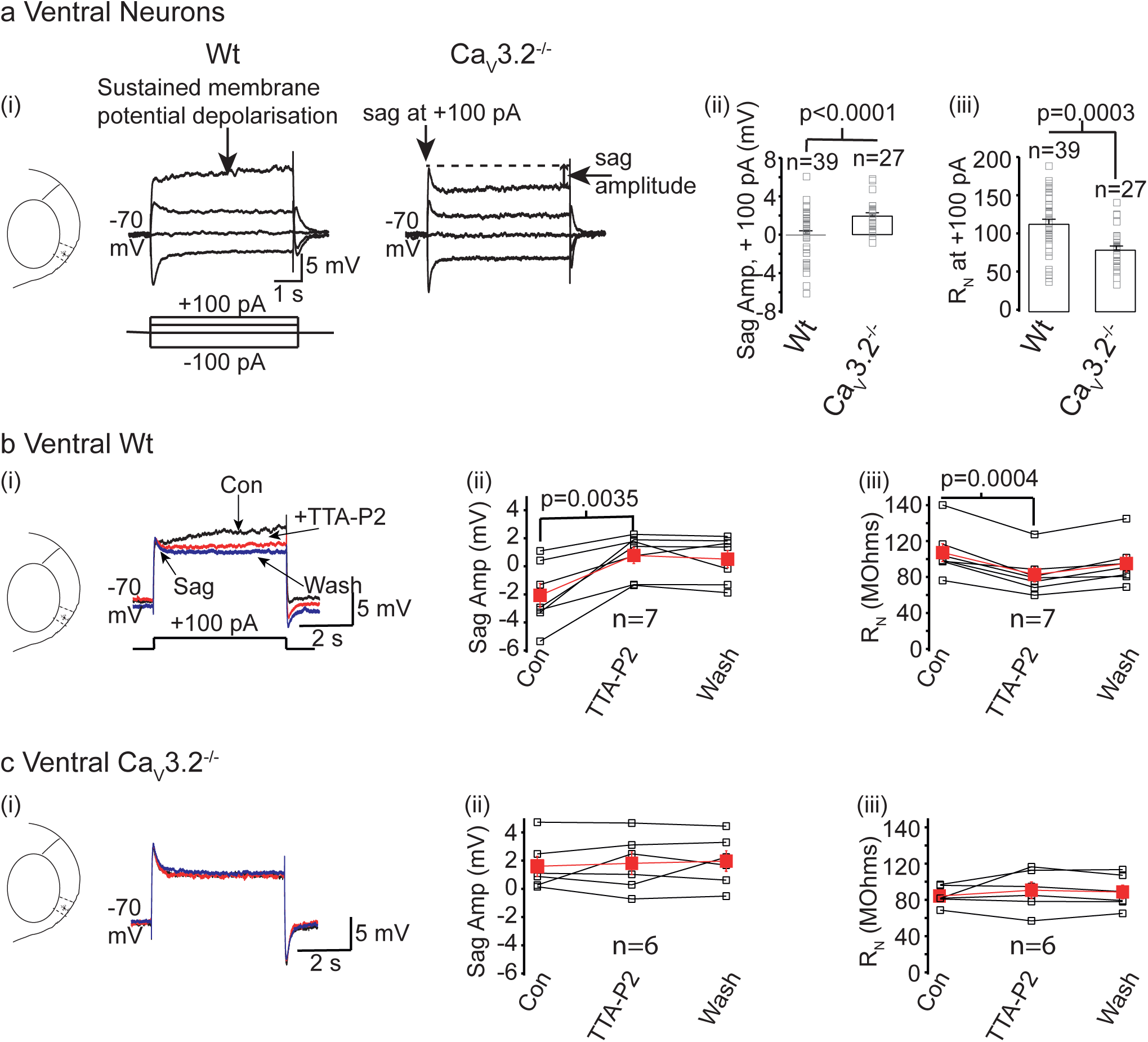
Ca_V_3.2 Ca^2+^ currents mask the membrane potential sag and enhance R_N_ at depolarizing potentials in ventral wildtype neurons. a(i) Example recordings obtained from ventral wildtype and Ca_V_3.2^-/-^ neurons in response to 5 s steps of -100 pA, +50 pA, +100 pA from a fixed potential of -70 mV. The sag amplitude was measured as the difference in voltage between the peak sag amplitude and the steady state voltage. The scale shown applies to both traces in the panel. **a(ii), a(iii)** Graphs showing the individual (open squares) and average (bars) sag amplitudes (amp) and R_N_ obtained with +100 pA current step respectively for ventral wildtype and Ca_V_3.2^-/-^ neurons. The numbers of observations for each group are indicated in parenthesis above the bars. Significance was determined using a two way ANOVA test corrected for multiple comparisons using a Bonferroni constant. The **b(i), c(i)** Representative traces showing the response to 5 s long, +100 pA current steps under control conditions (black), in the presence of TTA-P2 (red) and following washout of TTA- P2 (blue) in ventral wildtype and Ca_V_3.2^-/-^ neurons respectively. **b(ii), c(ii), b(iii), b(iii)** Graphs depicting the individual (black squares) and mean (filled red squares) values of the sag amplitude elicited using the +100 pA depolarizing step and the R_N_ measured using this step at a potential of -70 mV under control conditions, in the presence of TTA-P2 and after washout of TTA-P2. In all graphs, significance was determined using paired t-tests.

Stellate neurons express high densities of HCN channels (Garden et al., 2008; Giocomo and Hasselmo, 2008, 2009). These channels activate with hyperpolarization and de-activate with depolarization resulting in prominent membrane potential sags (Santoro and Shah, 2020). Indeed, as expected, hyperpolarizing pulses or small (+50 pA) subthreshold pulses resulted in membrane potential sags that were similar in amplitude in both ventral wildtype and Ca_V_3.2^-/-^ neurons (Supp Table 1) and indicates that HCN currents were unaffected by Ca_V_3.2 subunit knockdown in these neurons. Interestingly, though, the membrane potential sag due to large, +100 pA, subthreshold depolarizing pulses was significantly reduced in ventral wildtype neurons both at RMP and at -70 mV compared with Ca_V_3.2^-/-^ neurons (Fig 3a(i), Fig 3a(ii)). This implies that the initial voltage jump generated with large subthreshold depolarizing steps activated T-type, Ca_V_3.2 Ca^2+^ currents in ventral wildtype stellate neurons, instigating a prolonged membrane potential depolarisation and masking the sag (Fig 3a).

In support of the above notion, application of 100 nM TTA-P2 or 50 μM NiCl_2_ onto ventral wildtype, but not ventral Ca_V_3.2^-/-^, neurons resulted in a distinct membrane potential sag and reduction of R_N_ measured using depolarizing +100 pA potentials (Fig 3b, Fig 3c, Supp Fig 3). Neither of these compounds had any effect on the membrane potential sag or R_N_ measured using hyperpolarizing potentials in either ventral wildtype or Ca_V_3.2^-/-^ neurons (Supp Table 2). These findings reinforce the notion that Ca_V_3.2 Ca^2+^ channels are activated with subthreshold depolarizing potentials in ventral wildtype neurons, mask the membrane potential sag due to de-activating HCN channels and cause a sustained depolarization to enhance R_N_ and excitability.

### Ca_V_3.2 currents significantly influence the mEC stellate neuron R_N_ dorsal-ventral gradient at positive potentials

Next, we asked if Ca_V_3.2 currents affect R_N_ in dorsal neurons, even though these channels have little effect on their firing rates. We found no difference in dorsal wildtype and Ca_V_3.2^-/-^ neuron R_N_ or membrane potential sag measured with depolarizing or hyperpolarizing steps at either RMP or -70 mV (Fig 4a, Supp Table 1). Moreover, application of TTA-P2 or NiCl_2_ had little effect on dorsal wildtype or Ca_V_3.2^-/-^ neuron R_N_ or membrane potential sag elicited using depolarizing or hyperpolarizing steps at either RMP or -70 mV (Supp Fig 3, Supp Fig 4 Supp Table 2). Thus, these results further suggest that, unlike ventral neurons, dorsal neuron exhibit little or no Ca_V_3.2 currents.

**Fig 4:**
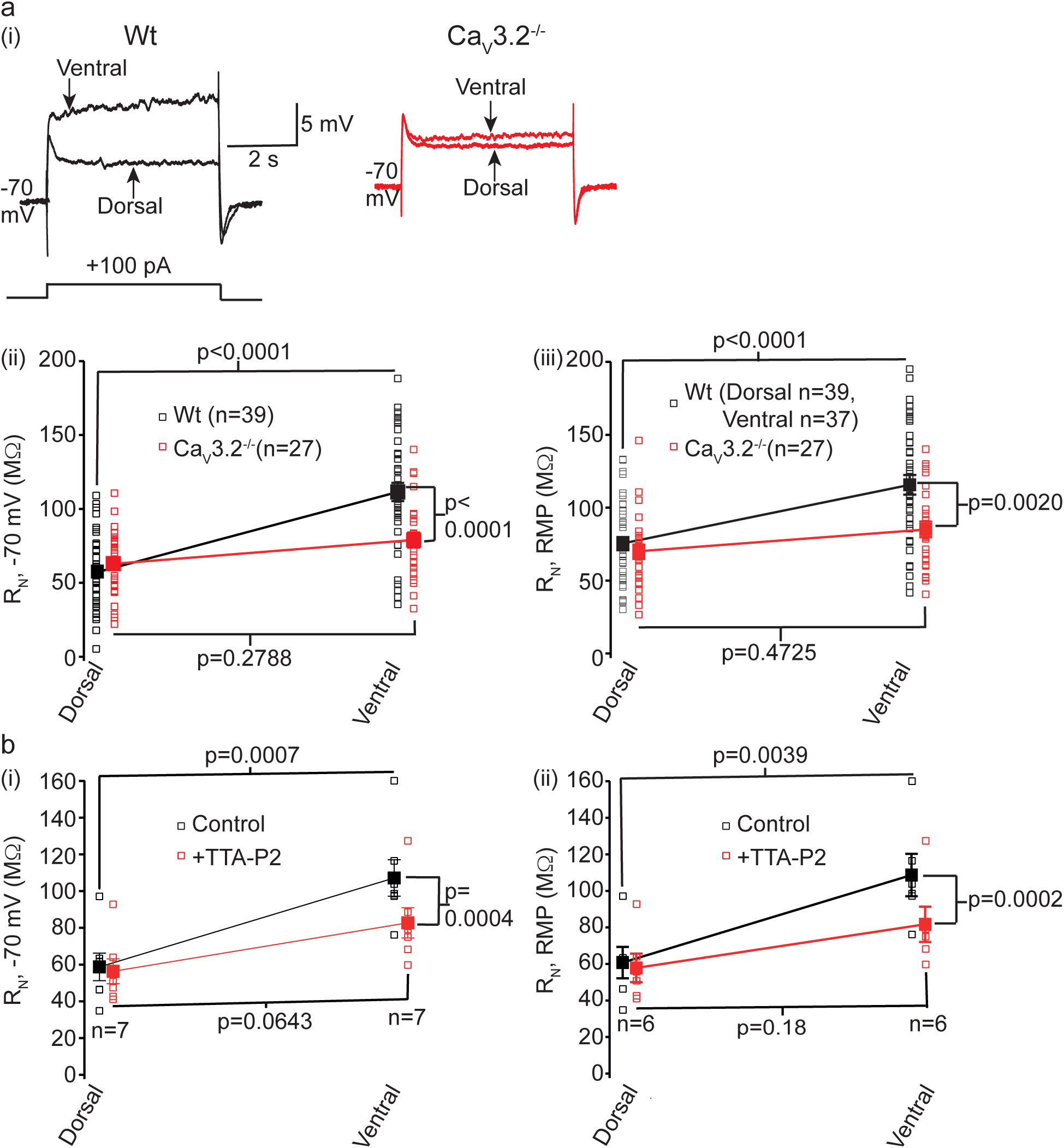
Ca_V_3 currents significantly influence the depolarizing R_N_ dorsal-ventral gradient in mEC stellate neurons. **a(i)** Representative traces from dorsal and ventral wildtype and Ca_V_3.2^-/-^ neurons when 5 s long, +100 pA depolarizing current steps were applied at a fixed potential of -70 mV. The scale shown applies to all traces. **a(ii), a(iii)** Graphs depicting the individual (open squares) and mean (closed squares) wildtype (black) and Ca_V_3.2^-/-^ (red) dorsal and ventral mEC stellate neuron R_N_ values obtained by applying 5 s long, +100 pA square pulses at -70 mV or RMP respectively. The numbers of observations for each group are indicated in parenthesis. **b(i), b(ii)** Plots illustrating the individual (open squares) and mean (closed squares) wildtype dorsal and ventral mEC stellate neuron R_N_ values obtained using 5 s, +100 pA current steps under control conditions (black) and in the presence of TTA- P2 (100 nM; red) at -70 mV or RMP respectively. The number of observations for each group are indicated on the graphs. Statistical significance for comparison of ventral and dorsal neurons was determined using a two-way ANOVA corrected for multiple comparisons using a Bonferroni constant. When determining statistical significance between ventral/dorsal groups with and without TTA-P2 application, paired t-tests were employed.

Consistent with the biophysical properties of Ca_V_3.2 currents, R_N_ measured using depolarizing 100 pA steps at -70 mV or at RMP was significantly greater in ventral than dorsal wildtype neurons (Supp Table 1, Fig 4a). This dorsal-ventral gradient in R_N_ was abolished in Ca_V_3.2^-/-^ stellate neurons (Fig 4a). Moreover, this dorsal-ventral R_N_ gradient was significantly reduced in the presence of TTA-P2 either at -70 mV or RMP (Fig 4b). This is primarily due to TTA-P2 causing a significant reduction in R_N_ measured using depolarizing pulses in ventral wildtype neurons (Fig 3, Fig 4b) but having little effect on this parameter in dorsal wildtype neurons (Supp Fig 4, Fig 4b). These findings indicate that by selectively enhancing R_N_ in ventral neurons, Ca_V_3.2 Ca^2+^ channels play a vital role in setting the dorsal-ventral gradient in R_N_ at depolarizing potentials and, therefore, action potential firing rates in mEC stellate neurons.

### Ca_V_3.2 Ca^2+^ currents are predominantly located in ventral wildtype stellate neurons

Since the loss of Ca_V_3.2 or application of T-type Ca^2+^ channel inhibitors affects ventral wildtype stellate neuron activity only, we investigated whether T-type, Ca_V_3.2 Ca^2+^ current density is greater in ventral wildtype stellate neurons compared with dorsal wildtype stellate neurons. We obtained whole-cell voltage-clamp recordings of T-type Ca^2+^ currents in the presence of N-type, P/Q-type, R-type, and L-type Ca^2+^ channel inhibitors as well as Na^+^ and K^+^ channel inhibitors (see Materials and Methods). T-type Ca^2+^ currents were activated and isolated using the protocol shown in Supp Fig 5a, which enables near-maximal activation of the current (Chemin et al., 2002). Interestingly, T-type Ca^2+^ currents were present at three times the density in ventral wildtype neurons compared with dorsal wildtype neurons (Fig 5a, 5b, 5c). Moreover, the T-type Ca^2+^ current amplitude in dorsal wildtype neurons was similar to that in dorsal Ca_V_3.2^-/-^ and ventral Ca_V_3.2^-/-^ neurons, implying that the dorsal wildtype neuron Ca_V_3.2 Ca^2+^ current density is minimal. Further, the small T-type Ca^2+^ currents present in dorsal and ventral Ca_V_3.2^-/-^ neurons indicate that Ca_V_3.1/Ca_V_3.3 subunits might also be expressed in these neurons. Nevertheless, these results robustly support the notion that T-type, Ca_V_3.2 Ca^2+^ currents are predominantly localised to ventral neurons only.

**Fig 5:**
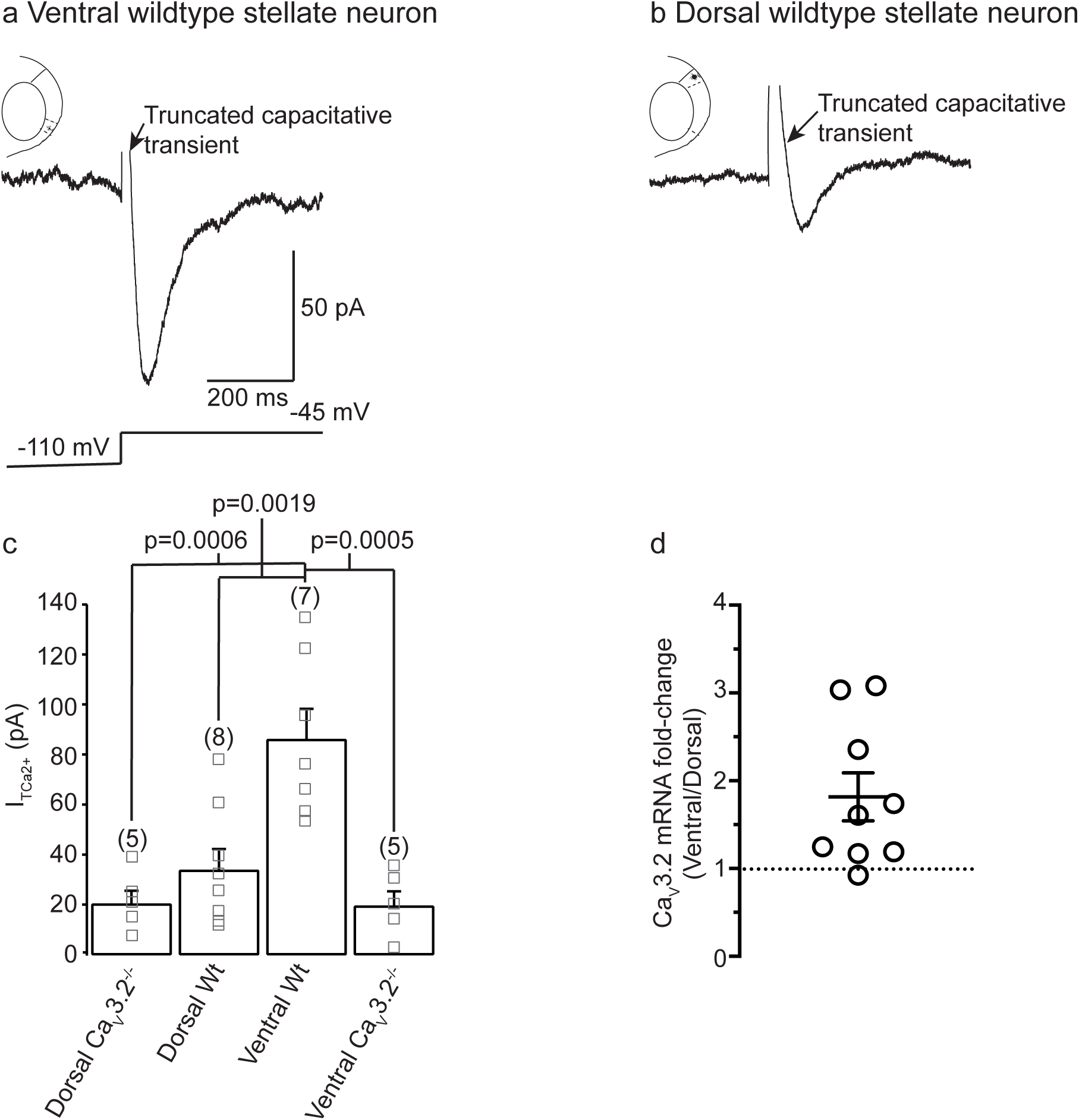
Ca_V_3 currents are present at significantly higher densities in ventral than dorsal wildtype neurons. **a, b** Typical absolute Ca_V_3 Ca^2+^ currents produced at the near maximal potential of -45 mV in ventral and dorsal wildtype stellate neurons respectively. The full protocol to generate the currents is illustrated in Supp Fig 5a. The scale associated with the recording in **a** applies to the trace shown in **b** too. **c** Bar graph showing the average (filled squares) and individual (open square) T-type Ca^2+^ current amplitudes obtained in dorsal Ca_V_3.2^-/-^, dorsal wildtype, ventral wildtype and ventral Ca_V_3.2^-/-^ stellate neurons. The numbers in parenthesis indicate the numbers of observations for each group of neurons. Statistical significance was determined using a two-way ANOVA corrected for multiple comparisons using a Bonferroni constant. **d** Graph depicting the fold change of Ca_V_3.2 mRNA in ventral, relative to dorsal mEC. Individual data (open circles) from 9 mice and mean ± SEM are shown (p = 0.0173, 1 sample 2 tailed test, compared to null hypothesis, dotted line shows no change).

The small T-type Ca^2+^ currents present in dorsal wildtype neurons precluded us from understanding their detailed biophysical characteristics including their activation and inactivation curves. The T-type Ca^2+^ currents in ventral wildtype neurons, though, were large enough to permit us to do this. In these neurons, the T-type Ca^2+^ currents had an average rise time constant of 11.9 ± 2.08 ms (n=7) and decay time constant of 77.5 ± 10.91 ms (n=7) at the peak voltage of -45 mV (Fig 5a, 5c, Supp Fig 5a, Supp Fig 5d). The activation and inactivation curves had half-maximal activation voltages (V_1/2_) of -53.9 ± 1.9 mV and -79.7 ± 1.3 mV respectively (Supp Fig 5a, Supp Fig 5b Supp Fig 5c). The slope (k) values of the activation and inactivation curves were 7.9 ± 1.0 mV^-1^ and 7.5 ± 0.9 mV^-1^ respectively (Supp Fig 5c). These V_1/2_ and k values are consistent with those reported for Ca_V_3.2 Ca^2+^ channels formed by expression in heterologous systems(Chemin et al., 2002), providing additional evidence that Ca_V_3.2 channels are likely to predominantly underlie the T-type Ca^2+^ current in ventral wildtype stellate neurons.

To further test the concept that Ca_V_3.2 expression differs between dorsal and ventral wildtype neurons, we performed quantitative reverse-transcriptase (RT)-PCR using micro- dissected adult mEC dorsal or ventral tissue (see Methods). The mRNA level for Ca_V_3.2 was greater in ventral compared to dorsal mEC in 8/9 WT mice examined. Using ΔΔC_T_ values, the fold-increase in ventral relative to dorsal tissue was 1.82 ± 0.27 (p = 0.0173, 1 sample t test, Fig 5d). These results further support the notion that ventral wildtype stellate neurons have a greater Ca_V_3.2-mediated T-type Ca^2+^ current than dorsal wildtype stellate neurons.

### T-type Ca^2+^ and subthreshold Na^+^ currents act together to enhance ventral wildtype stellate neuron excitability

The above findings show that T-type Ca^2+^ currents are very fast inactivating currents in mEC ventral stellate neurons (Fig 5a, Supp Fig 5a, Supp Fig 5d). This was surprising given that our experimental findings indicate that T-type, Ca_V_3.2 Ca^2+^ currents generate a sustained depolarization when subthreshold depolarizing steps are applied in ventral neurons (Fig 3).

This is critical for raising R_N_ (Fig 3, Fig 4) and action potential firing rates (Fig 1, Fig 2). This raises the question of how fast inactivating T-type Ca_V_3.2 Ca^2+^ currents might cause sustained depolarizations with subthreshold depolarizing stimuli in ventral neurons (Fig 1, Fig 2, Fig 3, Supp Fig 3a). To understand this, we generated a computational model of a ventral stellate neuron that incorporated known currents including an HCN current, K^+^ currents and a transient Na^+^ current (see Supp Table 4, Materials and Methods). We also integrated our experimental T-type Ca^2+^ current activation and inactivation characteristics and density in the cell soma only (Supp Table 4, Materials and Methods). Under these conditions, a subthreshold current pulse of +100 pA resulted only in a sag with no sustained depolarization (Supp Fig 6a). This suggests T-type Ca^2+^ currents on their own are unlikely to cause sustained depolarization in ventral neurons and indicates that T-type Ca^2+^ currents may act in concert with other subthreshold-activated voltage-gated channels to cause the sustained depolarisation in ventral neurons.

In addition to HCN currents, ventral stellate neurons possess axo-somatic subthreshold-activated persistent Na^+^ currents, albeit at a lower density than in dorsal wildtype neurons (Bant et al., 2020; Hargus et al., 2013; Magistretti and Alonso, 1999). Given this, we asked whether incorporation of this current together with somatic T-type Ca^2+^ currents in our computational model would reproduce the characteristic sustained depolarization produced with subthreshold current pulses in ventral wildtype neurons (Fig 2c, Fig 3a, Supp Fig 3a). Indeed, a +100 pA depolarizing subthreshold pulse now produced a prolonged depolarization with an undiscernible sag (Supp Fig 6b).

Next we tested whether T-type Ca^2+^ currents were a prerequisite for activating the persistent Na^+^ currents. Thus, we removed T-type Ca^2+^ currents from our model but left persistent Na^+^ currents intact. This resulted in a sag but no sustained depolarization with subthreshold current pulses (Supp Fig 6b). Hence, T-type Ca^2+^ currents are required to activate persistent Na^+^ currents to produce the sustained depolarization in ventral stellate neurons.

Given that the above strongly suggests that T-type Ca^2+^ and persistent Na^+^ currents together enhance R_N_, we examined whether T-type Ca^2+^ currents would modulate action potential firing in the presence of persistent Na^+^ currents in our model. In agreement with our experimental results (Fig 1, Fig 2, Supp Fig 3a), we found that removal of T-type Ca^2+^ currents reduced action potential numbers elicited by depolarizing pulses in our model (Supp Fig 6c). Hence, these findings provide additional evidence that T-type Ca^2+^ currents play a vital role in enhancing ventral wildtype stellate neuron action potential firing rates. To experimentally test the notion that T-type Ca^2+^ currents act in concert with persistent Na^+^ currents to cause sustained depolarizations with subthreshold pulses and thereby increase ventral wildtype stellate neuron activity, we applied riluzole (10 μM) followed by co-application of riluzole and the T-type Ca^2+^ current inhibitor, TTA-P2 (100 nM). Riluzole inhibits persistent Na^+^ currents more than resurgent Na^+^ currents(Bant et al., 2020). As expected, external application of riluzole inhibited action potential generation in both ventral wildtype and Ca_V_3.2^-/-^ stellate neurons significantly within 10 min (Fig 6a, Fig 6b). The resting membrane potential, as well as the R_N_ and sag due to a hyperpolarizing, -100 pA, 5 s long current pulse were similar before and after riluzole application in wildtype and Ca_V_3.2^-/-^ neurons (Supp Table 5). Interestingly, superfusion of riluzole, like T-type Ca^2+^ current inhibitors, resulted in a sag being revealed with depolarizing pulses and suppression of the sustained depolarization induced by +100 pA current pulses in ventral wildtype, but not Ca_V_3.2^-/-^, neurons (Fig 6a, 6b). Consequently, R_N_ was reduced in ventral wildtype stellate neurons but remained unchanged in ventral Ca_V_3.2^-/-^ neurons (Fig 6a). Subsequent co-application of riluzole and TTA-P2 for 20 min further enhanced the sag generated with the depolarizing +100 pA step in ventral wildtype stellate neurons only (Fig 6a). This, though, did not additionally alter R_N_ or the number of action potentials generated with depolarizing steps in these neurons (Fig 6a). Co-application of riluzole and TTA-P2, on the other hand, had little effect in ventral Ca_V_3.2^-/-^ stellate neurons (Fig 6b). Since riluzole either in the absence or presence of TTA-P2 had little effect on the R_N_ or sag produced by depolarizing steps in ventral Ca_V_3.2^-/-^ neurons, these findings support our hypothesis that the sag caused by deactivation of HCN channel currents activates T-type Ca^2+^ currents followed by Na^+^ currents in ventral wildtype stellate neurons.

**Fig 6:**
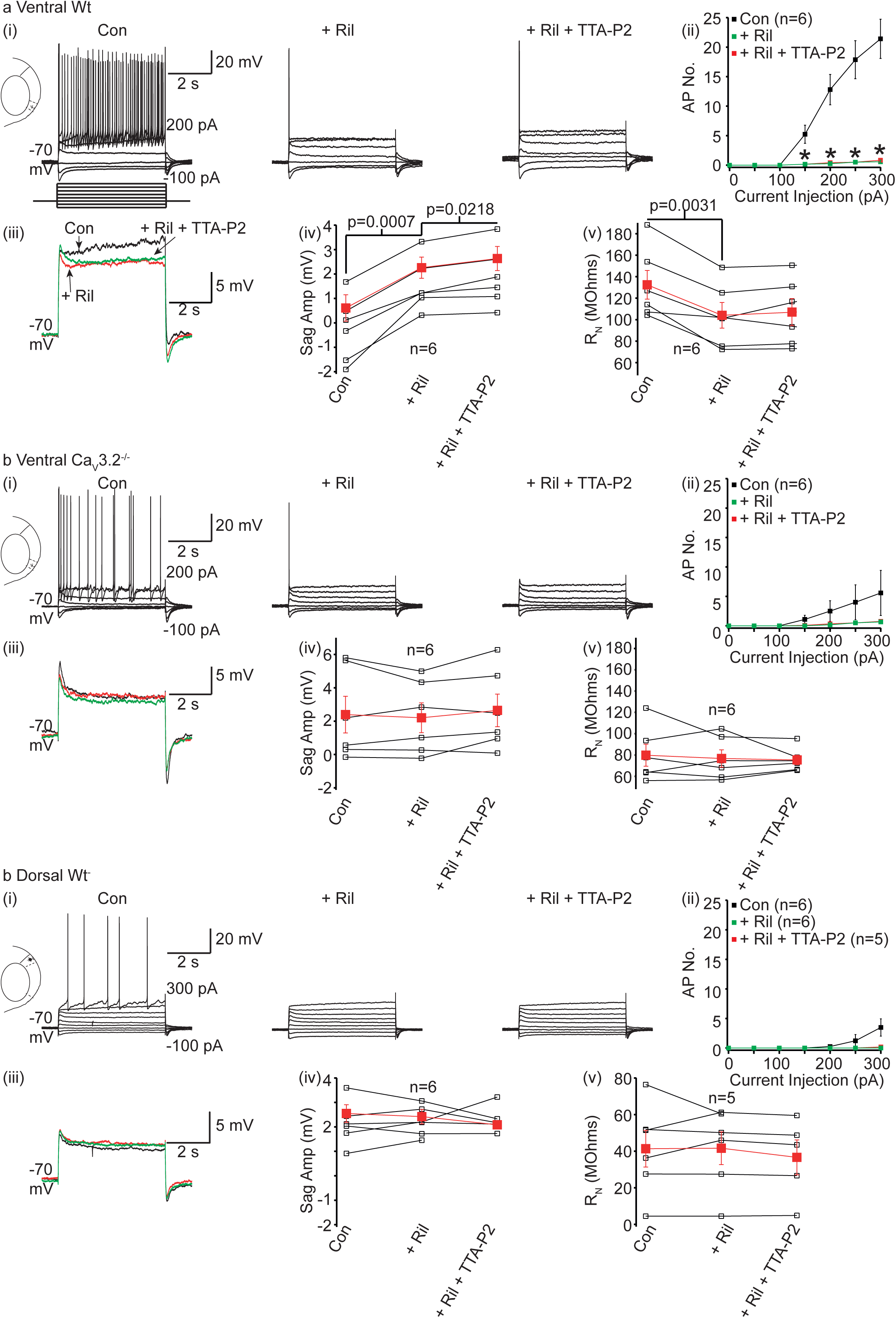
Ca_V_3 currents act together with subthreshold Na^+^ currents to enhance ventral wildtype stellate neuron R_N_ and excitability. **a(i), b(i), c(i)** Representative recordings from ventral wildtype, ventral Ca_V_3.2^-/-^ and dorsal wildtype stellate neurons respectively under control conditions, in the presence of riluzole (Ril, 10 μM) and subsequent co-application of riluzole and TTA-P2 (100 nM). The recordings were obtained at a potential of -70 mV by applying 5 s current pulses in 50 pA increments as shown in the schematic in **a**. The minimum and maximum current injections are indicated next to the first trace in each panel. The scale associated with the first trace in each panel applies to all traces within the panel. **a(ii), b(ii), c(ii)** Graphs depicting the mean number of action potentials elicited with depolarizing current pulses in the absence (con), presence of riluzole and subsequent co- application of riluzole and TTA-P2. The numbers of observations are indicated in parenthesis. Asterisks indicate significance at p<0.05 when action potential numbers under control conditions and in the presence of riluzole were compared using paired t-tests (exact p values are stated in Supp Table 7). **a(iii), b(iii), c(iii)** Superimposed typical traces obtained when a +100 pA current pulse was applied to ventral wildtype, ventral Ca_V_3.2^-/-^ and dorsal wildtype stellate neurons under control conditions (black), with riluzole (red) and following co-application of riluzole and TTA-P2 (green). **a(iv), a(v), b(iv), b(v), c(iv), c(v)** The individual (open squares) and mean (filled red square) sag amplitude (amp) and R_N_ values from ventral wildtype, ventral Ca_V_3.2^-/-^ and dorsal wildtype stellate neurons under control conditions, following application of riluzole alone and after subsequent co-application of riluzole and TTA-P2. Numbers of observations are indicated in brackets. NB for dorsal wildtype neurons, 6 individual recordings were obtained before and after application of riluzole. However, subsequent co-application of riluzole with TTA-P2 could only be recorded in 5/6 neurons. Paired t-tests were performed to determine if there was statistical significance, with those p values less than 0.05 being stated on the figure.

If T-type Ca^2+^ currents are required for the activation of persistent Na^+^ currents (as our experimental results and computational model suggest), then it can be hypothesized that riluzole will have little effect on R_N_ produced using depolarizing pulses in dorsal wildtype neurons which express high densities of subthreshold Na^+^ currents and HCN currents but have little T-type Ca^2+^ current (Fig 5b, 5c) (Bant et al., 2020; Garden et al., 2008; Giocomo and Hasselmo, 2008, 2009). Indeed, 10 min application of 10 μM riluzole had little effect on the R_N_ and sag due to depolarizing +100 pA step in dorsal wildtype stellate neurons (Fig 6c). Riluzole still significantly reduced action potential numbers (Fig 6c). Moreover, riluzole inhibited subthreshold oscillations in these neurons, consistent with previous reports (data not shown) (Klink and Alonso, 1993). As expected, the RMP as well as R_N_ and sag due to hyperpolarizing -100 pA current pulses were unaffected by riluzole in these neurons (Supp Table 5). Moreover, co-application of riluzole and TTA-P2 did not have any additional effects on the action potentials generated with depolarizing stimuli or on the R_N_ or sag induced by either depolarizing +100 pA or hyperpolarizing -100 pA steps (Fig 6c, Supp Table 5). These findings further support the notion that in the absence of T-type Ca^2+^ currents, the depolarization caused by the sag due to deactivating HCN currents by subthreshold depolarizing stimuli is by itself insufficient to activate subthreshold Na^+^ currents.

### Ca_V_3.2 channels prolong EPSP decay and integration in ventral wildtype mEC stellate neurons

*In vivo*, neuronal activity is strongly influenced by excitatory postsynaptic potentials (EPSPs) generation and integration. Thus, we investigated whether Ca_V_3.2 Ca^2+^ currents influenced EPSP amplitudes and decay in ventral stellate neurons and thereby modulate their activity *in vivo*. Ca_V_3.2 channels are also present presynaptically within the mEC and influence synaptic release under certain conditions (Huang et al., 2011). Thus, to avoid the additional complexity caused by possible differences in endogenous synaptic release onto neurons in wildtype and Ca_V_3.2^-/-^ tissue, we generated EPSPs by injecting current waveforms using a defined α function in the presence of glutamate and GABA receptor inhibitors (see Materials and Methods; known as αEPSPs). The rise time of αEPSPs was set to be 1 ms under control conditions. This protocol resulted in αEPSPs of similar amplitude in ventral wildtype and Ca_V_3.2^-/-^ stellate neurons (Fig 7a(i)). The αEPSP decay time constant, though, was significantly reduced in ventral Ca_V_3.2^-/-^ neurons compared with wildtypes (Fig 7a(ii)). Application of 100 nM TTA-P2 also reproduced the reduction in αEPSP decay time constants in ventral wildtype neurons whilst having little effect on their amplitude (Fig 7b(i), Supp Table 6). As expected with the reduced density of T-type Ca^2+^ currents in dorsal neurons, Ca_V_3.2 loss or inhibition of these currents with TTA-P2 had little effect on αEPSP amplitude or decay in these neurons (Fig 7a(ii), Fig 7b(ii)). Hence, these findings indicate that Ca_V_3.2 channels are activated during EPSPs and prolong their delay in ventral wildtype stellate neurons.

**Fig 7:**
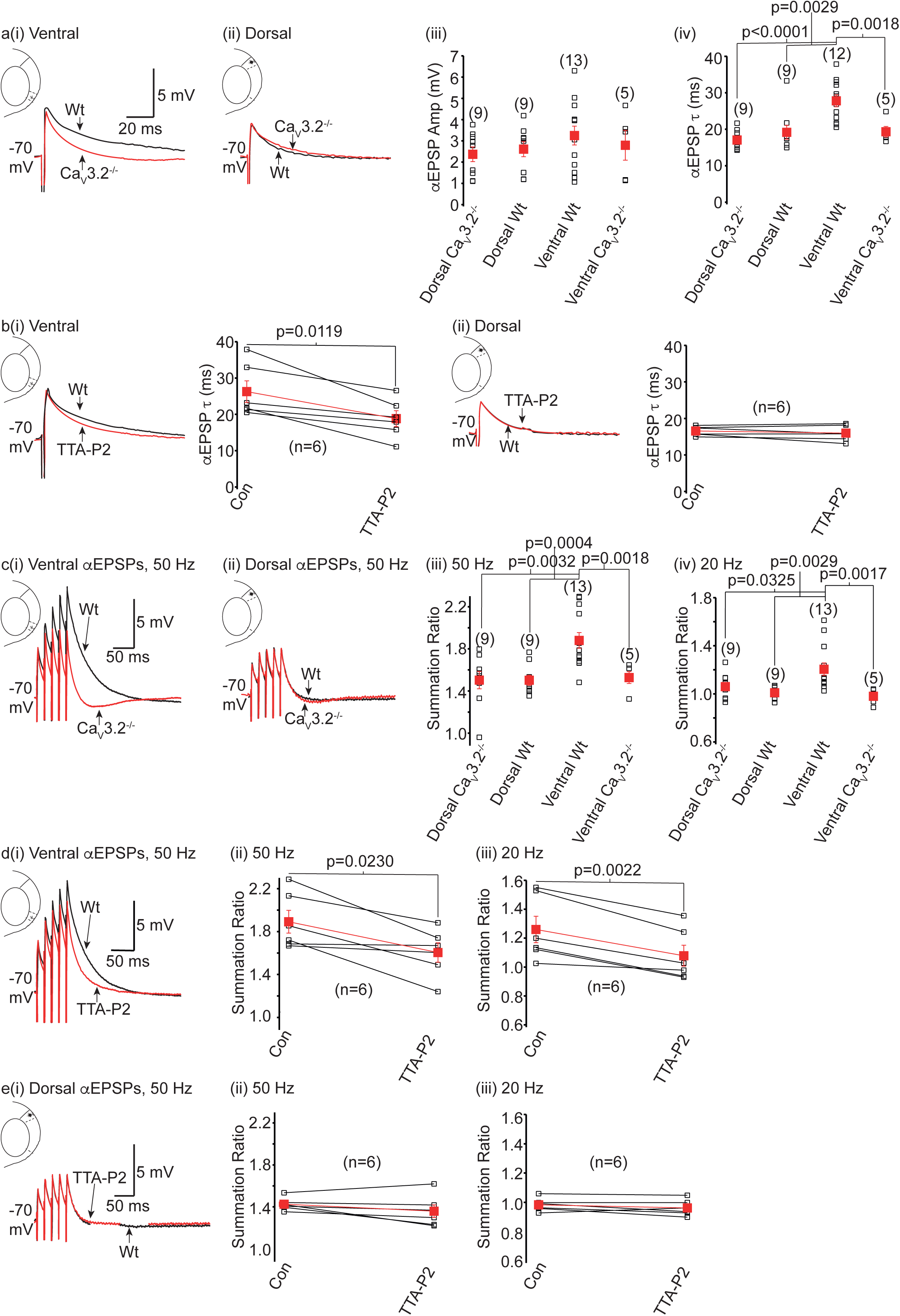
Ca_V_3.2 Ca^2+^ currents regulate *α*EPSP decay time constants and summation in ventral wildtype stellate neurons alone. a(i), a(ii) Single αEPSP recordings in ventral and dorsal stellate neurons. Black and red traces represent recordings from wildtype (Wt) and Ca_V_3.2^-/-^ neurons respectively. The scale applies to **a(i)** and **a(ii)**. **a(iii), a(iv)** Graphs depicting individual (open black squares) and mean (red filled squares) αEPSP amplitudes and decay time constants (t) at -70 mV in all four groups. Significance values were determined using two-way ANOVA with Fisher’s Least Significance Difference (LSD) post hoc test. **b(i), b(ii)** Representative traces showing single αEPSPs in ventral and dorsal wildtype stellate neurons respectively in the absence (black) and presence (red) of TTA-P2. The graphs illustrate the individual (open squares) and average (red filled squares) single αEPSP decay time constant (τ) values under control conditions and following 20 min application of TTA-P2. Paired t-tests were performed to determine significance between control values and those in the presence of TTA-P2. **c(i), c(ii)** Example recordings of 5 αEPSPs at 50 Hz in ventral and dorsal stellate neurons respectively obtained from Wt and Ca_V_3.2^-/-^ mice. **c(iii), c(iv)** Plots showing the summation ratios obtained from trains of 5 αEPSPs evoked at 50 Hz and 20 Hz respectively in dorsal Ca_V_3.2^-/-^, dorsal wildtype, ventral wildtype and ventral Ca_V_3.2^-/-^ stellate neurons. Significance values were determined using two-way ANOVA with Fisher’s Least Significance Difference (LSD) post hoc test**. d(i), e(i)** Representative traces of 5 αEPSPs elicited at 50 Hz before (black trace) and after 20 min TTA-P2 application (red trace) in ventral and dorsal wildtype stellate neurons respectively. **d(ii), d(iii), e(ii), e(iii)** Graphs displaying the individual (open black square) and mean (red filled square) summation ratio values obtained from trains of 50 Hz and 20 Hz αEPSPs in the absence (con) and presence of TTA-P2 in ventral and dorsal wildtype stellate neurons. Paired t-tests between control EPSP summation ratios and EPSP summation ratios in the presence of TTA-P2 were performed to determine significance. In all graphs, the numbers of observations for each group are shown in parenthesis.

The decay time constant strongly influences EPSP integration and thereby action potential generation *in vivo*. To test whether Ca_V_3.2 Ca^2+^ currents affect EPSP integration, we generated trains of 5 αEPSPs elicited at either 20 Hz or 50 Hz. We found that Ca_V_3.2 channel abolition resulted in reduced summation of αEPSP trains elicited at either 50 Hz or 20 Hz in ventral, not dorsal, stellate neurons (Fig 7c) Consistent with this, 20 min external application of TTA-P2 onto ventral, but not dorsal, wildtype stellate neurons reduced the summation of a train of 5 αEPSPs at either 20 Hz or 50 Hz (Fig 7d, 7e). These findings, therefore, robustly support the notion that T-type Ca^2+^ currents are activated during EPSPs, and influence their integration and signal processing *in vivo* in ventral wildtype stellate neurons.

## Discussion

Here, we have provided substantial evidence to demonstrate that ventral, but not dorsal, mEC layer II stellate neurons possess T-type Ca^2+^ currents which play a fundamental role in shaping their intrinsic excitability and processing synaptic information. We found that the T-type Ca^2+^ currents encoded by Ca_V_3.2 α1 subunits were predominantly present in ventral stellate neurons (Fig 5c). These T-type Ca^2+^ currents were activated with depolarizing stimuli in these neurons and contributed significantly to enhancing R_N_ as measured with long depolarizing steps in these neurons (Fig 3, Fig 4). This was unexpected since these currents have fast inactivating kinetics (Fig 5a; Supp Fig 5). We showed that this effect was due to T-type Ca^2+^ currents triggering the activation of subthreshold-activated persistent Na^+^ currents (Fig 6, Supp Fig 5). Indeed, even though dorsal stellate neurons have larger densities of persistent Na^+^ currents compared with ventral stellate neurons, equivalent depolarizing pulses were unable to activate these currents to increase R_N_ (Fig 6c). Hence, T-type Ca^2+^ currents are critical for activating persistent Na^+^ currents and augmenting R_N_ in mEC ventral stellate neurons. As a consequence of the significantly larger R_N_ in ventral stellate neurons relative to dorsal stellate neurons, there was a greater probability of action potentials being generated with depolarizing stimuli and enhanced EPSP summation in ventral stellate neurons (Fig 1, Fig 2, Fig 7, Supp Fig 3). Thus, our findings reveal a unique mechanism by which ventral stellate neurons regulate their intrinsic and synaptic activity. Further, importantly, our results indicate that ventral and dorsal neurons express distinct voltage-dependent conductances that differentially influence their excitability.

### A novel mechanism by which T-type Ca^2+^ channels exert their effects in neurons

Unusually, we found that T-type Ca^2+^ currents activated persistent Na^+^ currents to enhance ventral wildtype stellate neuron R_N_ and action potential firing propensity (Fig 6, Supp Fig 6). This mechanism has not previously been described in other cell types yet. In most neurons, T-type Ca^2+^ channels are activated during the rebound potential following a hyperpolarizing pulse, and facilitate action potential generation with rebound potentials (Zamponi et al., 2015). Further, in so-called ‘pacemaker’ neurons such as corticothalamic neurons, T-type Ca^2+^ currents have been shown to act with HCN channels to generate oscillations and rhythmic spiking. Whilst there was a rebound potential following a hyperpolarizing pulse in ventral wildtype neurons, the amplitude of this was similar irrespective of the absence or presence of Ca_V_3.2 Ca^2+^ currents (data not shown) and no action potentials were generated with rebound potentials. Moreover, similar proportions of ventral wildtype and Ca_V_3.2^-/-^ neurons had intrinsic oscillations, indicating that T-type Ca^2+^ currents do not contribute to the generation of these oscillations in ventral wildtype neurons. Hence, in ventral wildtype stellate neurons, T-type Ca^2+^ currents are predominantly activated with depolarizing potentials, which led to persistent Na^+^ current activation to cause sustained depolarization and enhanced neuronal excitability.

The interaction between T-type Ca^2+^ channels and persistent Na^+^ channels in ventral wildtype stellate neurons is likely to be voltage-dependent as this effect was reproducible in a simple computational model in which the persistent Na^+^ current biophysical characteristics are not altered following T-type Ca^2+^ current activation (Supp Fig 5). The possibility, though, that Ca^2+^ entry via T-type Ca^2+^ channels may also rapidly and reversibly phosphorylate Na^+^ channels to enhance the persistent Na^+^ current in ventral mEC layer II stellate neurons cannot be excluded. Certainly Ca^2+^-dependent modification of Na_V_1.6 channels, which principally encode for persistent Na^+^ currents in neurons (Hargus et al., 2013; Katz et al., 2018), increased the persistent Na^+^ current in hippocampal CA1 pyramidal neuronal dendrites (Yu et al., 2018) and cerebellar Purkinje neurons (Zybura et al., 2020). Moreover, Ca^2+^- calmodulin dependent kinase, CaMKII, phosphorylates Na_V_1.2 channels, that predominantly underlie transient Na^+^ currents, to generate a persistent Na^+^ current in heterologous systems (Thompson et al., 2017). It should be noted, however, that there is very little T-type Ca^2+^ current active at RMP in ventral mEC layer II stellate neurons (Fig 1, Fig 2,, Supp Fig 3, Supp Table 1, Supp Table 2). Hence, it is unlikely that Ca^2+^ entry via these channels at rest will tonically activate CaMKII and thus generate persistent Na^+^ currents in these neurons. Therefore, though we cannot rule out the possibility that Ca^2+^-dependent phosphorylation of Na^+^ channels might be a mechanism for transiently boosting the persistent Na^+^ current during depolarizing stimuli, our results strongly indicate that T-type Ca^2+^ currents and persistent Na^+^ currents act in concert in a voltage-dependent manner to enhance ventral mEC layer II stellate neuron R_N_ and activity (Fig 6, Supp Fig 5).

Whilst our work shows that that peri-somatic T-type Ca^2+^ channels interact with axo-somatic persistent Na^+^ currents to influence ventral stellate R_N_ and excitability, this interaction may also be present in other neuronal compartments such as axons and dendrites as these channels have been shown to be localized here too in other neurons including within the mEC (Evans et al., 2017; Huang et al., 2011; Isope et al., 2012; Leresche and Lambert, 2017; Magee et al., 1995; Martinello et al., 2015; Zamponi et al., 2015). Our data indicated that T-type Ca^2+^ channels have little effect on the action potential threshold *per se* in stellate neurons (Supp Fig 2, Supp Table 3), indicating that if these channels are present in axons, they do not affect stellate neuron intrinsic excitability. Whilst our computational model suggests that somatic T-type Ca^2+^ channels alone are sufficient to reproduce our experimental findings (Supp Fig 5, Supp Table 5), we cannot ignore the possibility that they may also be in dendrites, akin to data from hippocampal, substantia nigra pars compacta, corticothalamic and Purkinje neurons (Evans et al., 2017; Isope et al., 2012; Leresche and Lambert, 2017; Magee et al., 1995; Martinello et al., 2015). In corticothalamic and Purkinje neurons, these channels critically regulate synaptic potential properties and, consequently, synaptic plasticity (Isope et al., 2012; Leresche and Lambert, 2017). Since mEC stellate neurons have active dendrites that contain, at least, voltage-gated Na^+^ channels (Schmidt-Hieber et al., 2017), it is plausible that T-type Ca^2+^ channels might also be expressed in mEC stellate neuron dendrites and influence synaptic potential shapes and integration, as we find in the soma (Fig 7).

Moreover, since persistent Na^+^ channels can also be localized to dendrites (Yu et al., 2018), a voltage-dependent interaction between these channels and T-type Ca^2+^ channels might also exist in dendrites. This might serve to amplify and lengthen EPSPs and boost their integration, thereby facilitating their propagation to the soma to influence action potential generation.

### T-type Ca^2+^ currents as means for autonomous regulation of ventral hippocampal circuit activity

Recent evidence indicates that, like in the hippocampus, ventral and dorsal mEC circuit connections are distinct and have different functions (Cembrowski and Spruston, 2019; Fanselow and Dong, 2010; Steffenach et al., 2005; Strange et al., 2014). As yet, there is little information on how these circuits might be regulated individually to enable them to perform their functions independently. The dominant presence of T-type Ca^2+^ currents in ventral mEC stellate neurons might represent a physiological mechanism for selectively manipulating ventral mEC circuit activity. Since Ca_V_3.2 Ca^2+^ channel activity can be modulated by diverse neurotransmitters such as acetylcholine and dopamine (Bender et al., 2012; Evans et al., 2017; Jin et al., 2019; Martinello et al., 2015; Zamponi et al., 2015), alterations in T-type Ca^2+^ current activity might represent a unique mechanism by which ventral hippocampal circuit activity and functions can be selectively regulated.

In summary, we have shown that ventral mEC stellate neurons have enhanced intrinsic excitability compared with dorsal mEC stellate neurons. This is mediated by the predominant presence of T-type Ca^2+^ currents in ventral mEC stellate neurons. Whilst we have focused on understanding the mechanisms underlying ventral mEC stellate neuron intrinsic excitability, our findings might also be applicable to other brain regions. Indeed, ventral hippocampal dentate gyrus and CA1 neurons also exhibit increased excitability compared with their dorsal counterparts (Papatheodoropoulos, 2018). As with mEC stellate neurons, this difference between ventral and dorsal hippocampal neurons has thus far been attributed to altered density of HCN/K^+^ channels across the ventral/dorsal axis (Dougherty et al., 2013; Malik et al., 2016; Malik and Johnston, 2017). Ventral hippocampal neurons might, though, express distinct active conductances from dorsal neurons, which might influence their activity, akin to ventral mEC stellate neurons. Hence, distinct expression profiles of voltage-gated conductances, even within particular regions of the brain, are likely to have diverse effects on neuronal excitability and, thereby, influence neural circuit activity and function.

## Acknowledgments

This work was supported by a European Research Council Starter Independent Grant (GA 260725 IRPHRCSTP, M.M.S) and a UCL IMPACT studentship (A.T, M.M.S, A.C.D). This research was also funded in part by the Wellcome Trust (Grant No. 206279\Z\17\Z; A.C.D). For the purpose of open access, the author has applied a CC BY public copyright license to any author accepted manuscript version arising from this submission. In addition, we thank Mr. S. Martin (Biosciences, UCL) for genotyping services, Ms K. Chan (UCL SoP) for assisting with immunohistochemistry and morphological analysis and Dr. T. Evans (UCL Institute of Neurology, UCL) for initial analysis of subthreshold oscillation frequency and discussions.

## Author Contributions

All electrophysiological experiments and analysis was performed by AT and MMS. EG and MM generated the computational model and carried out the computational modelling. WSP and ACD did the RT PCR experiments and analysis. MMS wrote the manuscript with contributions from ACD and MM.

## Materials and Methods

### Ethics and Experimental Approval

All animal experiments were approved by the UCL animal welfare and ethics review body and were performed with approved UK Home Office personal and project licenses. All animals were housed under 12 hr dark/light cycles and were provided with *ad libitum* food and water. Ca_V_3.2 heterozygotes (Ca_V_3.2^+/-^) breeding pairs were used to generate wildtype and Ca_V_3.2 null (Ca_V_3.2^-/-^) littermates for experiments as previously described(Chen et al., 2003; Huang et al., 2011).

### Acute slice preparation

5-8 week old wildtype and Ca_V_3.2^-/-^ mice were terminally anaesthetized using a ketamine/xylazine mixture. Intra-cardiac perfusion was then carried out using an ice-cold modified artificial cerebral spinal fluid (ACSF) of the following composition (mM): 2.5 KCl, 1.25 NaH_2_PO_4_, 25 NaHCO_3_, 0.5 CaCl_2_, 7 MgCl_2_, 7 dextrose, 110 choline chloride. 300 μm thick parasagittal slices were obtained using an LT 1200S vibratome (Leica Microsystems, UK). Slices containing the mEC were identified and selected as previously described (Pastoll et al., 2012b). These slices were transferred to a chamber maintained at 35 °C and containing ACSF (mM): 125 NaCl, 2.5 KCl, 1.25 NaH_2_PO_4_, 25 NaHCO_3_, 2 CaCl_2_, 2 MgCl_2_, 10 dextrose, pH 7.3 maintained with 95 % O_2_/ 5 % CO_2_ mixture. Slices were stored at 35 °C for 25 min, after which they were kept at room temperature for 40 min.

### Electrophysiological recordings

For patch-clamp recordings, the slices were transferred to a submerged chamber perfused at 3-5 ml/min with ACSF supplemented with 10 μM CNQX, 50 μM DL-AP5, 10 μM bicuculline and 1 μM CGP 55845 and maintained at 32-36 °C. The dorsal and ventral mEC were initially identified under low magnification (x10) using an Olympus BX5141 microscope (Microscope Service and Sales, UK). The parasubicular/dorsal mEC border could be visualized using low magnification objectives under a microscope (Olympus BX51W1) equipped with differential infrared contrast optics. The ventral mEC border was estimated from position of the CA1/subicular border(Pastoll et al., 2012b). Stellate neurons located within 30 % of the mEC dorsal border was classified as dorsal (Fig 1a). Correspondingly, those situated within 30 % of the ventral border were categorized as ventral (Fig 1b).

#### Current-clamp recordings

Whole-cell current-clamp recordings were obtained from visually-identified stellate neurons present in either dorsal or ventral regions of the slice. Patch pipettes had a resistance of 5-8 MΩ when filled with the following solution (mM): 120 KMeSO4, 15 KCl, 10 HEPES, 0.2 EGTA, 2 MgCl2, 4 Na2ATP, 0.3 Tris-GTP, 14 Tris – phosphocreatinine, 0.2% neurobiotin, pH adjusted to 7.3 with 1 M KOH (295 mOsm/l). Recordings were made using an Axoclamp 200B amplifier (Molecular Devices Ltd) and acquired using pClamp 10.4 (Molecular Devices Ltd). Data were filtered at 10 kHz and sampled at 50 kHz. 5 s long, hyperpolarizing and depolarizing square pulses were injected every 10 s either at the resting membrane potential or at a fixed potential of -70 mV to determine the intrinsic membrane characteristics and action potentials elicited in response to depolarizing stimuli. To record action potential characteristics, short 10 ms square pulses were applied. αEPSPs were generated using the equation:

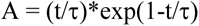

where A and τ represent the amplitude and rise time constant respectively. τ was set to be 1 ms. In all whole-cell current-clamp experiments, series resistance was in the order of 10 – 20 MΩ and recordings were discarded if this changed by > 20 %.

#### Voltage-clamp recordings

Whole-cell voltage-clamp recordings were obtained stellate neuron somata. The ACSF was supplemented with 1 μM tetrodotoxin, 2 mM CsCl_2_, 10 mM tetraethylammonium chloride, 0.1 mM 4-aminopyradine, 20 μM nifedipine, 0.2 μM ω- agatoxin IVA, 0.2 μM ω-conotoxin GVIA, 0.2 μM SNX-482 and maintained at 32- 36 °C. Patch pipettes were filled with an intracellular solution consisting of (mM): 120 CsCl_2_, 1 CaCl_2_, 5 MgCl_2_, 10 EGTA, 10 HEPES, 4 Na_2_ATP, 0.3 Tris-GTP, 14 Tris – phosphocreatinine, 0.2 % neurobiotin and pH adjusted to 7.3 with 1 mM CsOH (295-300 mOsm/l). Recordings were obtained using an Axopatch 200 B (Molecular Devices Ltd) and acquired using pClamp 10.4 (Molecular Devices Ltd). Data were filtered at 10 kHz and sampled at 50 kHz. Cells were voltage-clamped at -70 mV. To activate the T-type Ca^2+^ current, a 1s pre-pulse to -100 mV followed by 1 s pulses ranging from -90 mV to -45 mV were applied every 10 s (see Supp Fig 5a). The inactivation protocol consisted of applying 1 s pulses from -100 mV to -55 mV followed by a 1 s step to -50 mV every 10 s (Chemin et al., 2002) (Supp Fig 5b). A leak subtraction step consisting of a 1s step to -100 mV followed by a 1s step to -110 mV was applied between each protocol. Each protocol was repeated at least three times in the absence and after 20 min application of 100 nM TTA-P2 (Supp Fig 5).

### Neurobiotin Staining

Following electrophysiological recordings, all slices were fixed in 0.4 % paraformaldehyde solution for a minimum of 20 min followed by a three separate washes with phosphate buffered solution (PBS). The slices were then stained with Alexa Fluor 488 streptavidin conjugate antibodies (0.2 % in PBS) using a standard protocol (Huang et al., 2012). The slices were then mounted on microscope slides and stored at least overnight at 4 °C. The slices were subsequently viewed using a confocal microscope (Zeiss LSM 710) to confirm if the neuron recorded from was located in the dorsal or ventral mEC.

### Data Analysis

#### Whole-cell current clamp recording analysis

Clampfit 10.4 (Molecular Devices Ltd) was used. Sag amplitude was measured as the difference between the peak amplitude and the steady state potential for a given current pulse (Fig 2a). R_N_ was calculated by dividing the difference in steady state voltage in the last 25 ms of a 5 s +100 pA or -100 pA step at -70 mV with the current applied. Action potentials elicited by 5 s depolarizing steps were counted. The latency to spike was computed as the time for the first action potential to be initiated at the smallest depolarizing step applied.

To measure the action potential threshold, a phase plane plot was constructed (Supp Fig 2)(Martinello et al., 2015). For this, the voltage was differentiated with respect to time (dV/dt) and plotted against voltage. The threshold was defined as the voltage at the point of deflection for dV/dt to be greater than zero. The spike amplitude was the voltage difference between the threshold and the peak. The action potential half-width was then estimated at half the spike amplitude.

The αEPSP decay time constant (τ) was estimated by fitting the decay phase with a single exponential:

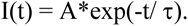

where I is the current amplitude, A is the peak amplitude, τ is the decay time constant (Huang et al., 2009). The summation ratio for a train of 5 αEPSPs at either 20 Hz or 50 Hz was the ratio of the amplitude of the 5^th^ αEPSP and the 1^st^ αEPSP.

*T-type Ca^2+^ current measurements* Currents generated by the activation and inactivation protocols were first leak subtracted in Clampfit 10.4. The resulting currents in the absence of TTA-P2 were then subtracted from those obtained in the presence of TTA-P2 to obtain isolated T-type Ca^2+^ currents. The peak amplitudes of these were measured at all voltages. To generate the activation and inactivation curve, the peak amplitudes at different voltages were expressed as a ratio of the maximal current amplitude produced by the protocol at any given voltage (I/I_Max_). This ratio was then plotted against the voltage (Supp Fig 4c). The curves were then fitted using Origin Pro 10 (OriginLab) with a Boltzmann equation:

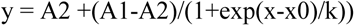

where A1 and A2 are the initial and final y values, x0 represents the half-maximal voltage (V_1/2_) and k is the slope of the curve.

In addition, the rise and decay time constants were estimated by fitting the currents obtained with activation protocol with a single exponential function as described. These were plotted against voltages and fitted in Origin Pro 10 using a second order polynomial function of the order:

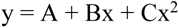

where A is an offset value and B and C are coefficients.

#### Sholl analysis, cell soma size measurements and estimation of cell location

Acquired confocal images were uploaded into Image J and z-stacks generated to obtain the full soma and dendritic arbor (Supp Fig 1). The total mEC length was measured from the parasubicular border to the ventral border. 30 % of this was calculated and cells located within the distance from either the dorsal or ventral borders were classified as dorsal or ventral respectively.

The cell soma area was estimated using Image J macros. For cells that had full-length dendrites extending to the edge of the slice, Sholl analysis was carried as previously described (Huang et al., 2012; Martinello et al., 2015). Concentric circles that were 10 μm apart were constructed around the soma. The number of dendrites crossing each circle were counted for each cell (Supp Fig 1b).

#### Statistical analysis

Group data are expressed as mean ± SEM. Sample size was based on our previous experience for doing similar type of electrophysiology experiments. For initial experiments for Fig 1, Fig 2, Fig 3, Fig 4, Supp Fig 3 and Supp Fig 4, the experimenter was blind to the mouse genotype until data analysis had been completed. To determine statistical significance at p < 0.05 between the four groups, a two-way ANOVA with Fisher’s Least Significance Difference (LSD) post hoc test were performed using GraphPad Prism 8.0. In some cases, a two-way ANOVA adjusted for multiple comparisons using a Bonferroni Constant was performed to avoid false positives, as specified in the figure legends. To determine if there were significant differences between control conditions and following application of a T-type Ca^2+^ channel inhibitor, paired t-tests were used. Significance (p values) are stated on figures or Supp Table 7.

### Materials and methods for slice preparation, electrophysiological analysis and neurobiotin staining

All salts with the exceptions of NaCl, KCl, KMeSO4 and PBS were purchased from Sigma Aldrich UK. These were acquired from Fisher Scientific UK. Neurobiotin was obtained from Vector Laboratories UK whilst DL-AP5, bicuculline CNQX, CGP 55845, tetrodotoxin, tetraethylammonium chloride, 4-aminopyridine and nifedipine were purchased from Abcam Ltd (UK). ω-agatoxin IVA, ω-conotoxin GVIA and SNX482 were acquired from Tebu Bio UK. Alexa 488 steptavidin conjugated antibodies were obtained from Life Technologies UK (now known as Thermo Fisher Scientific UK). TTA-P2 was a generous gift from Merck Laboratories USA.

### Quantitative PCR

400 μm mEC parasagittal slices were prepared from 6 - 8 week old wildtype mice as described above. The dorsal and ventral mEC layer II/III was micro-dissected from the slices and immediately frozen on dry ice. Tissue samples were disrupted using a rotor-stator homogenizer (Disperser T10, IKA, Staufen, Germany). Total RNA was extracted from the tissue using the RNeasy Lipid Tissue Kit (Qiagen) including an on-column DNase step. Reverse transcription (RT) was performed on matched amounts of RNA from Dorsal and Ventral samples from the same animal, using High capacity RNA-to-cDNA kit (Applied Biosystems). RT-negative controls were included. TaqMan qRT-PCR (40 cycles) performed with an Applied Biosystems 7500/7500 Fast Real-Time PCR system was used to determine the relative abundance of the Ca_V_3.2 α1 subunit in pair-matched samples (between 17 and 50 ng cDNA). The following TaqMan assays with TaqMan Gene Expression Master Mix were used in accordance to the manufacturer’s protocol (gene name: assay ID): Hypoxanthine phosphoribosyltransferase (*Hprt1)*: Mm00446968_m1, cacna1h (Ca_V_3.2): Mm00445382_m1. Measurements were performed in triplicate on independent RNA preparations from nine mice, and Ca_V_3.2 mRNA levels were normalised for expression of *Hprt1* mRNA. The experimenter was blind to the mouse genotype until data analysis had been completed.

Sample size was determined by previous experience of similar experiments. Comparison between dorsal and ventral tissue from each mouse was made by calculating ΔΔC_T_.

### Computational Modelling

All simulations were carried out using the NEURON simulation environment (v7.7.2) (Hines and Carnevale, 1997). For all simulations we used a reconstructed morphology of a stellate cell (courtesy of Prof. M. F. Nolan (University of Edinburgh, UK)(Garden et al., 2008)), with uniform passive properties (Cm = 1 μF/cm^2^, *R*m = 50kΩ/cm^2^, *R*a = 150 Ω cm). Temperature was set at 34°C. Active properties included a transient Na^+^ conductance, two types of K^+^ currents (a delayed rectifier and a Ca^2+^-dependent K^+^ conductance), a non-specific I_h_ current, a Ca^2+^ conductance modelling T-type Ca^2+^ currents, and a simple Ca^2+^-extrusion mechanism with a 500 ms time constant. Kinetics for the Ca^2+^-dependent K^+^ current was taken from a previously published model (ModelDB accession no. <u>112546</u>) (Shah et al., 2008). Parameters for the other currents are reported in Supp Table 4. All model and simulation files will be uploaded to the ModelDB database (https://senselab.med.yale.edu/modeldb/266797). Because the scope of this model was to test a specific hypothesis on the role of the interaction between T-type Ca^2+^ and persistent Na^+^ currents, rather than implement a full model for these cells, a more extensive parameter search was not performed. The peak conductance for all channels was manually adjusted following a trial-and-error procedure to reproduce the main effects observed in the experimental traces at 100 and 150pA. The effects of TTA-P2 application were modelled with a complete block of the T-type Ca^2+^ conductance.

---

**Supp Fig 1:**
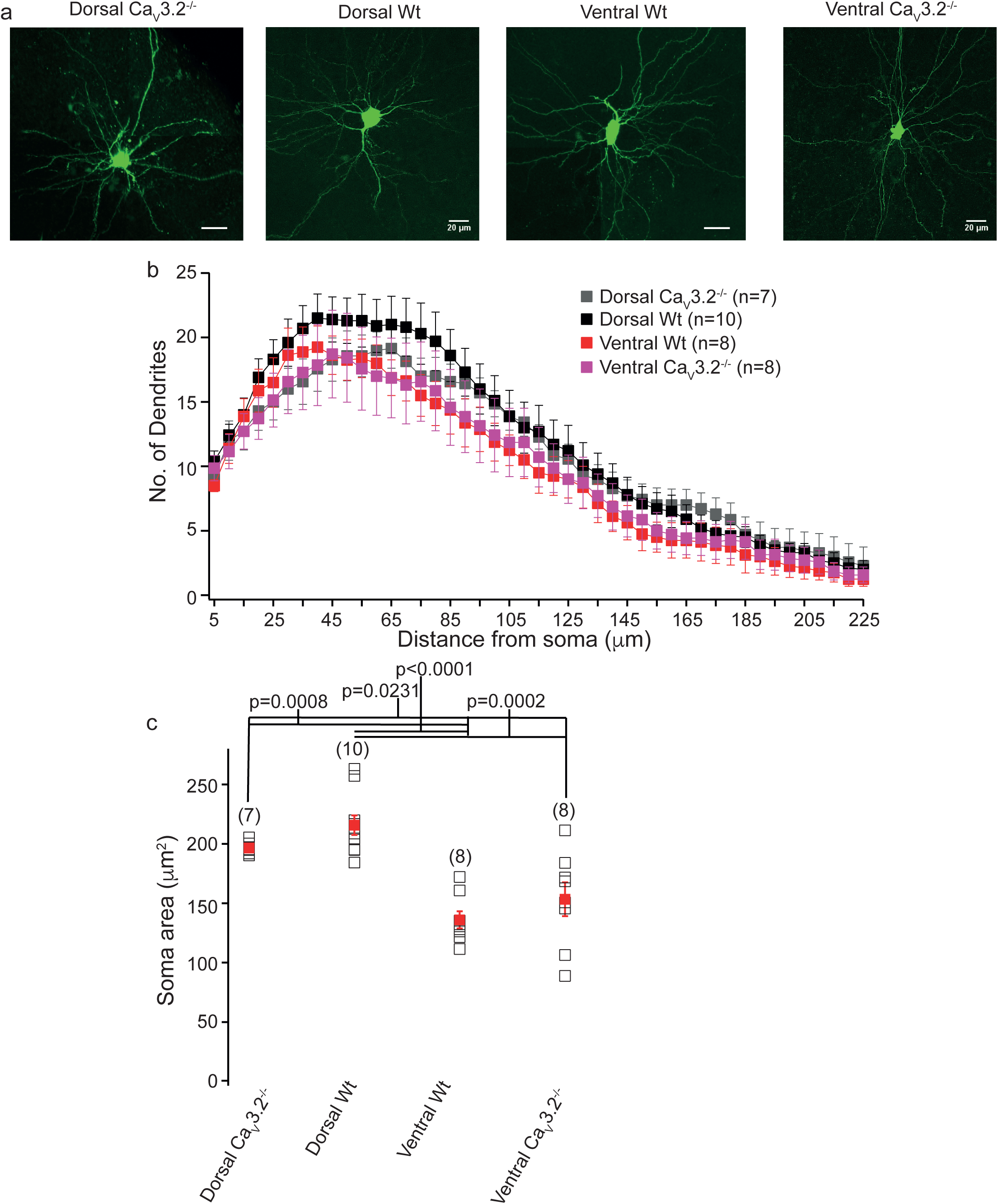
Morphological characteristics of wildtype and Ca 3.2^-/-^ dorsal and ventral mEC layer II stellate neurons. **a** Confocal images of wildtype (Wt) and Ca 3.2^-/-^ dorsal and ventral mEC layer II stellate neurons. These were filled with neurobiotin and stained using Alexa Fluor 488 (see **Methods**). The scale bar shown on each image corresponds to 20 μm**. b** Sholl analyis showing the number of dendrites imaged at a given distance from the soma in wildtype and Ca 3.2^-/-^ dorsal and ventral neurons. Each symbol represents the mean and SEM. **C**. Soma size of dorsal and ventral mEC layer II stellate neurons. Open (black) and filled (red) squares depict individual and mean data points respectively. The numbers of observations are in parenthesis. Asterisks indicate significance at p < 0.05 (two way ANOVA adjusted for multiple comparisons using a Bonferroni constant).

**Supp Fig 2:**
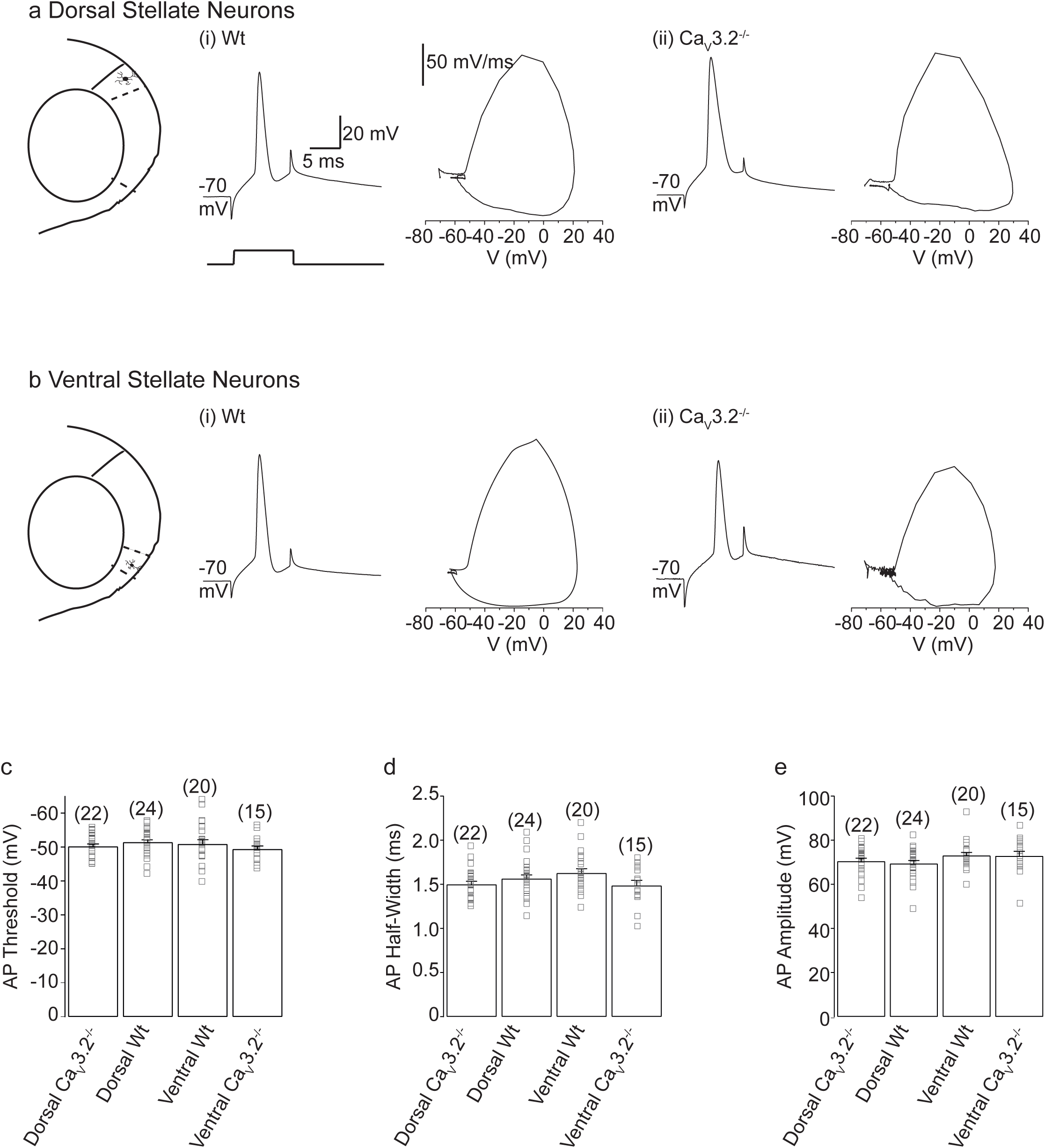
Ca 3.2 Ca^2+^ channels have little effect on action potential characteristics. **a, b** Example action potential traces from dorsal and ventral wildtype (**a(i), b(i)**) and Ca_V_3.2 null (**a(ii), b(ii)**) stellate neurons. Individual spikes were elicited using a 10 ms step from a fixed potential of -70 mV. The phase plane plots for the individual action potentials are shown next to the traces. The action potential threshold was measured from these. The scale bars associated with the single action potential and phase plane plot in **a(i)** applies to all traces and plots in **a** and **b**. **c, d, e** Bar graphs depicting the mean and SEMs for action potential threshold, half-width and action potential for all 4 groups of neurons respectively. Open squares represent individual values of the measurements for each group. The number of observations per group are shown in parenthesis above each bar. No significance at p < 0.05 was detected (statistics performed using a two-way ANOVA adjusted for multiple comparisons using a Bonferroni Constant).

**Supp Fig 3:**
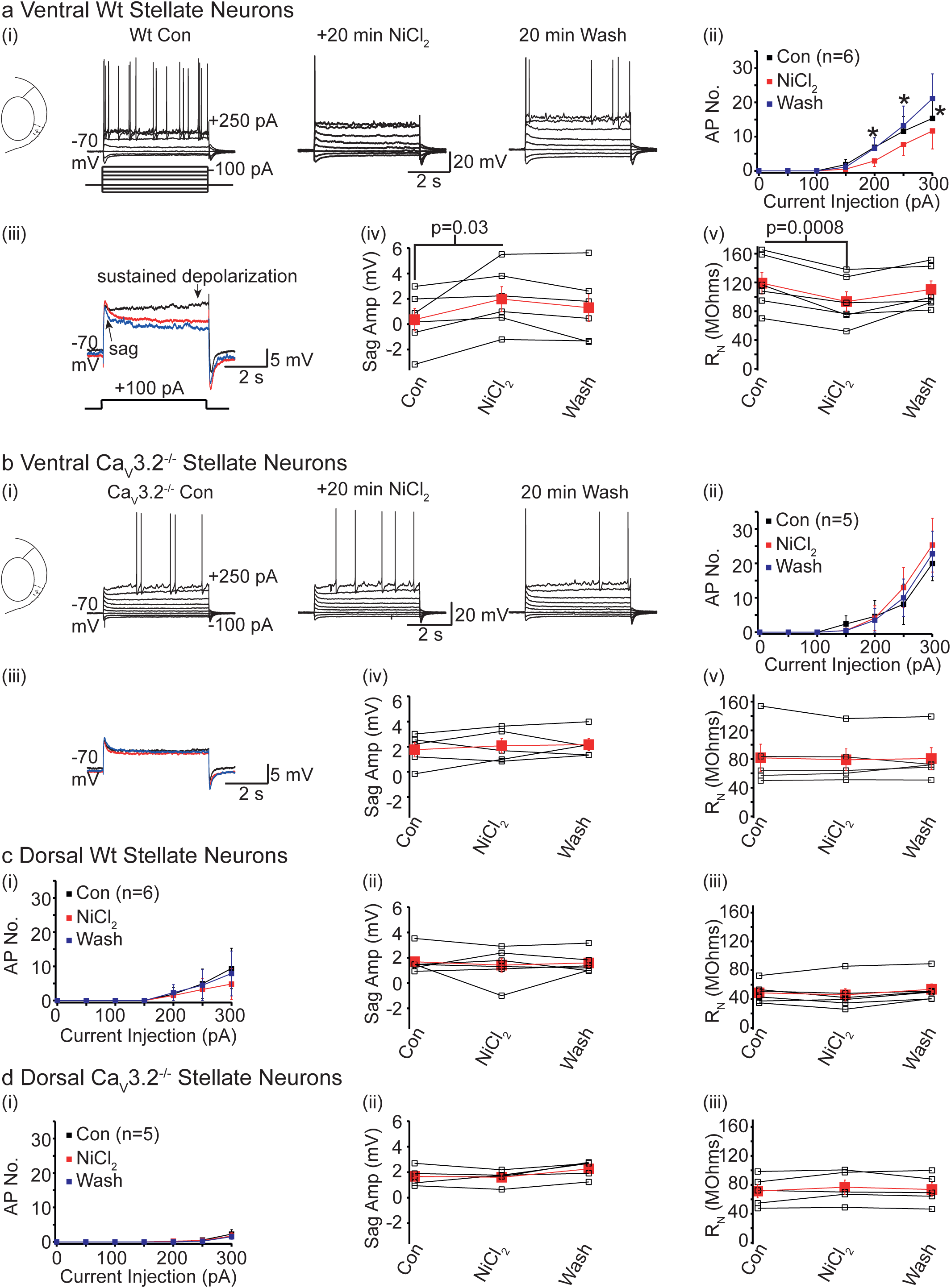
Ventral wildtype stellate neuron activity, depolarizing sag and input resistance at positive potentials is selectively affected by NiCl_2_. **a(i), b(i)** Representative whole-cell current clamp recordings from ventral wildtype and Ca 3.2^-/-^ stellate neurons in the absence (con), after 20 min of NiCl_2_ application and following 20 min washout respectively. The recordings were produced by applying 5 s long steps from -100 pA to +250 pA as shown in the schematic below the trace in **a(i)**. The scale shown in **a(i)** and **b(i)** applies to all traces within these panels. **a(ii), b(ii)** Graphs depicting the number of action potentials elicited by depolarizing steps in ventral wildtype and Ca 3.2^-/-^ stellate neurons respectively. The numbers of observations are in parenthesis. Asterisks signify significance at p< 0.05 when action potential numbers are compared under control conditions and in the presence of NiCl_2_ using paired t-tests (exact p values are stated in Supp Table 7). **a(iii), b(iii)** The +100 pA traces from the recordings shown in **a(i) and b(i)** respectively on an expanded time scale. The traces have been superimposed. **a(iv), b(iv), a(v), b(v)** Plots illustrating the sag amplitude and input resistance measured using the +100 pA current pulse without NiCl_2_, with NiCl_2_ and following washout of NiCl_2_ in ventral wildtype and Ca 3.2^-/-^ stellate neurons. Open black and filled red squares represent individual and mean values respectively. Control values are compared with those obtained after 20 min treatment with NiCl_2_ using paired t-tests. The p values are stated for when a significance of p < 0.05 was detected. **c, d** Graphs depicting action potential numbers with incremental 5 s, depolarizing steps **(i)**, sag amplitude with the +100 pA step **(ii)** and the input resistance obtained using the +100 pA step **(iii)** in dorsal wildtype and Ca_V_3.2^-/-^ stellate neurons respectively in the absence, presence and following washout of NiCl_2_. Filled squares with error bars represent mean and SEM values for all graphs while open squares depict individual values (**(ii), (iii)**).

**Supp Fig 4:**
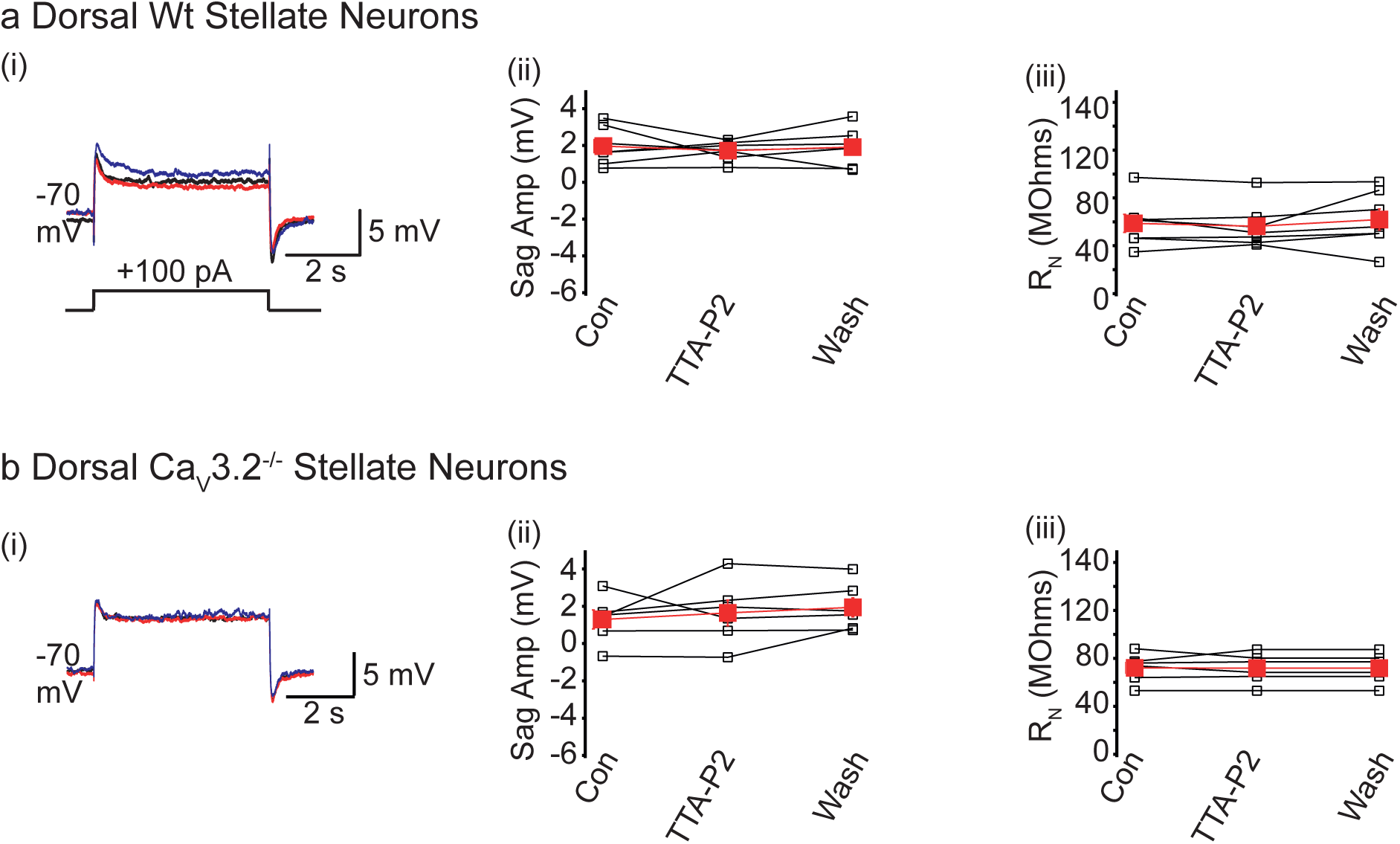
The T-type Ca^2+^ channel inhibitor, TTA-P2, has little effect on the sag or R_N_ in dorsal stellate neurons. **a(i), b(i)** Representative traces when 5 s long, +100 pA depolarizing steps were applied to dorsal wildtype (Wt) or Ca 3.2^-/-^ neurons under control conditions (black), in the presence of TTA-P2 (100 nM; red) and following washout (Wash, blue). **a(ii), b(ii), a(iii), b(iii)** Graphs depicting the sag amplitude and input resistance (R_N_) measured using this protocol under these conditions.

**Supp Fig 5:**
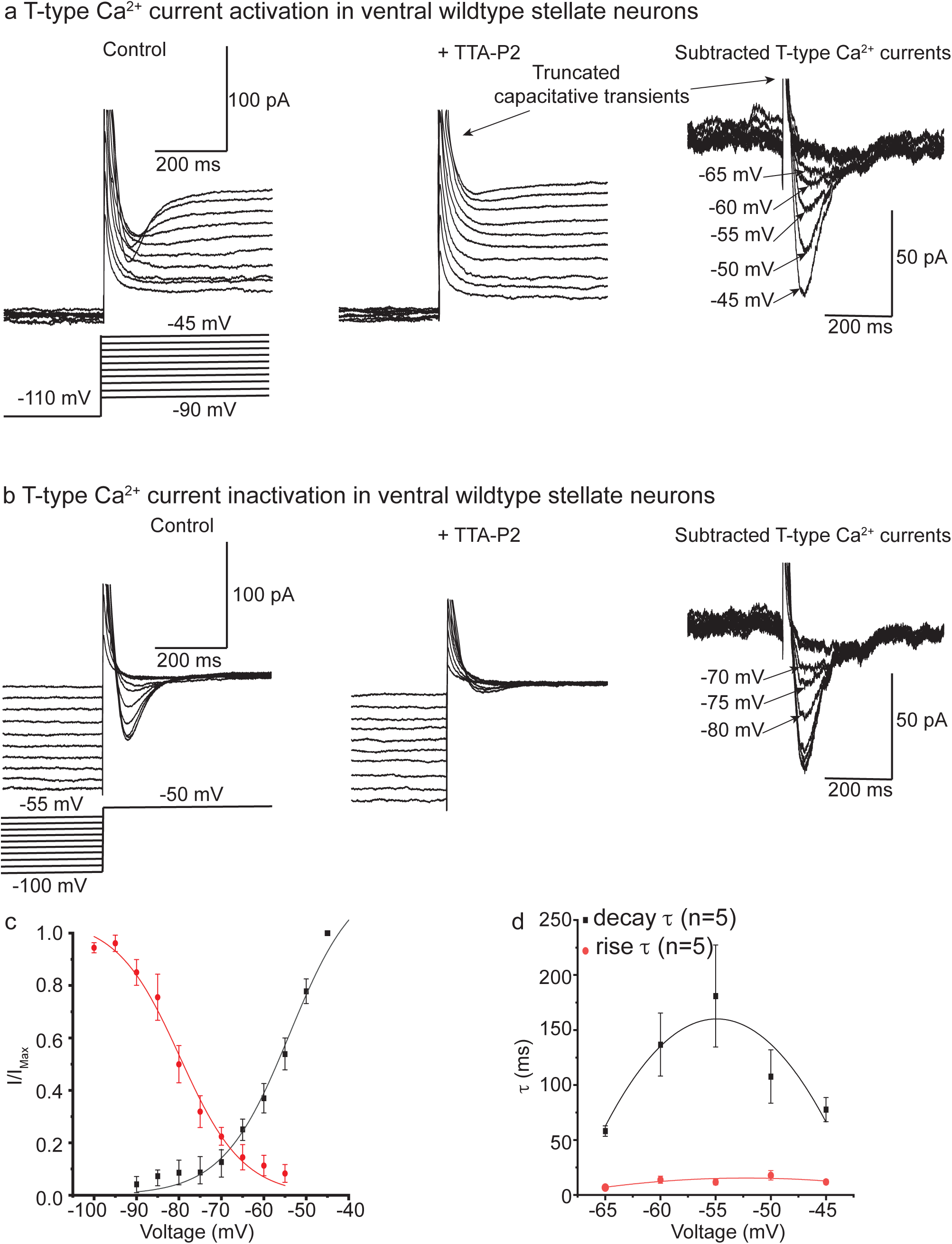
Biophysical properties of T-type Ca^2+^ currents in ventral wildtype stellate neurons. **a. b** Example raw traces of currents generated by the voltage protocols shown undr control conditions and in the presence of 100 nM TTA-P2. The scale bar associated with the control trace also applies to the trace recorded with TTA-P2. All traces have been leak-subtracted (see Methods). Recordings following 20 min TTA-P2 application were subtracted from those obtained under control conditions to isolate the T-type Ca^2+^ current (far right panels). Current amplitudes, decay time constants (τ) and rise τ values were obtained from these traces. **c** The activation (black) and inactivation (red) curve of T-type Ca^2+^ currents recorded from 7 ventral wildtype stellate neurons. Each point represents the mean and SEM. The values were fitted with a Boltzmann function (see Methods). **d** A plot showing the variance of the decay and rise τ with voltage. Each point represents the mean and SEM recorded from 5 different neurons.

**Supp Fig 6:**
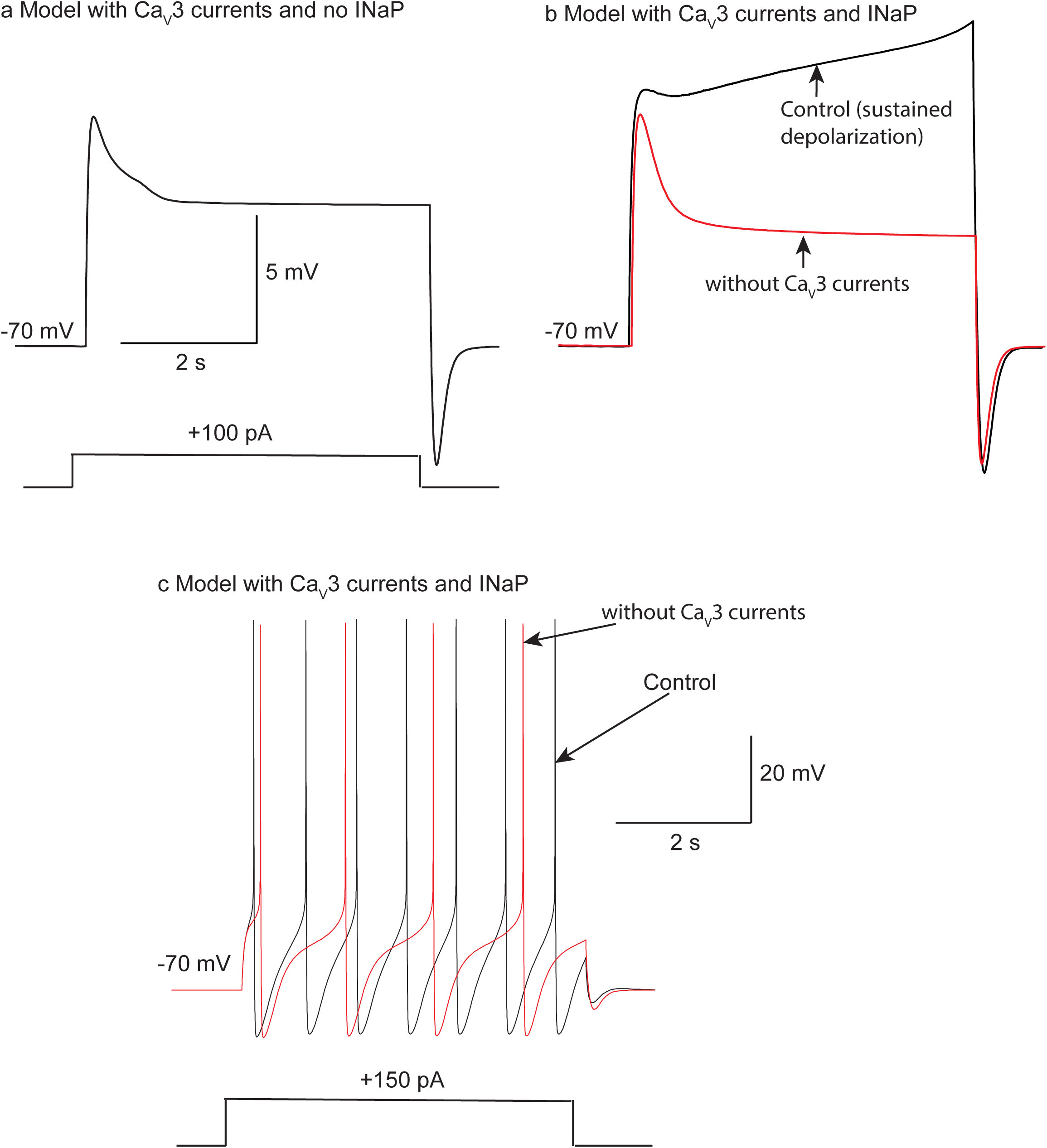
Computational model indicates that Ca 3 currents together with persistent Na^+^ currents boost excitability in ventral wildtype stellate neurons. **a** Simulations indicate that a computational model incorporating Ca 3 currents but no persistent Na^+^ currents (INaP) generates a sag only in response to a +100 pA subthreshold current pulse. **b** Simulation suggesting that when both Ca_V_3 and INaP currents were included (control; black trace), subthreshold +100 pA current pulses generated a sustained depolarization. Removal of Ca_V_3 currents (red trace) then resulted in a sag. **c** Simulations when larger suprathreshold currents were injected in a model containing both Ca_V_3 and INaP currents. The control (black) trace resulted in several action potentials. The number of action potentials was reduced when Ca_V_3 currents were removed (red trace).

